# Dysregulation of locus-specific repetitive elements in TCGA pan-cancers

**DOI:** 10.1101/2024.10.20.619138

**Authors:** Chao Wang, Chun Liang

## Abstract

Understanding the role of repetitive elements (REs) in cancer development is crucial for identifying novel biomarkers and therapeutic targets. This study investigated the locus-specific dysregulation of REs, including differential expression and methylation of REs, across 12 TCGA cancer types stratified by their genomic context (*i.e.*, genic and intergenic REs). We found uniquely dysregulated genic REs co-regulated with their corresponding transcripts and associated with distinct biological functions in different cancer types. Uniquely dysregulated intergenic REs were identified in each cancer type and used to cluster different sample types. Recurrently dysregulated REs were identified in several cancer types, with genes associated with up-regulated genic REs involved in cell cycle processes and those associated with down-regulated REs involved in the extracellular matrix. Interestingly, 4 out of 5 REs consistently down-regulated in all 12 cancer types were located in the intronic region of the TMEM252, a recently discovered tumor suppressor gene. TMEM252 expression was also down-regulated in 10 of 12 cancer types, suggesting its potential importance across a wide range of cancer types. With the corresponding DNA methylation array data, we found a higher prevalence of hypo-methylated REs in most cancer types (10 out of 12). Despite the slight overlaps between differentially expressed REs and differentially methylated REs, we showed that methylation of locus-specific REs negatively correlates with their expression in some cancer types.

## Introduction

### Repetitive elements (REs) and their regulations in the human genome

Repetitive elements (REs) are the most abundant type of sequences in the human genome^1^ can be classified into satellites (or tandem repeats) and transposable elements (TEs), with the latter further subdivided into RNA transposable elements and DNA transposable elements. REs are typically hierarchically classified with an increasing granularity from class, family, and subfamily (corresponding to repClass, repFamily, and repName hierarchies defined by the human Repeatmasker^2^) to locus-specific elements based on sequence similarities^3^. As arrays of repeated nucleotides, satellites can be further classified based on the increasing size of the repeated unit into microsatellites, minisatellites, satellites, and macrosatellites^4^. Microsatellites, also known as simple sequence repeat (SSR) or short tandem repeat (STR), are small (2-6 bp in repeat unit) tandem repeat accounting for about 3% of the human genome (most of the telomeric region are occupied by microsatellites)^4^. Compared to microsatellites, minisatellites are larger (about 15 bp in repeat unit) but fewer in the human genome (found in euchromatic regions of the genome). Due to their highly variable array size they are also known as variable number of tandem repeats (VNTRs). Most satellites are composed of either simple repeats or complex repeats with increased repeat unit length and complexity, mainly appearing at centromeres, pericentromeric regions and subtelomeric regions (*e.g.*, alpha-satellites are the most abundant satellite in human genome mainly occupying the centromere with a repeat unit of 171bps, accounting for half of the total satellite DNA; the human pericentromeric regions contains satellite I (17 bps and 25 bps), satellite II (10-80 bps), satellite III (5 bps and 10 bps) and alpha-satellite (68 bps and 220 bps)). The repeat unit in macrosatellite can be up to few kilobases in length (*e.g.*, human macrosatellites D4Z4 is 3.3 kb located at the subtelomeric region in chromosomes 4q35 and 10q26)^4^. While the tandem repeats are primarily found in centromeres and telomeres, TEs and their evolutionary relics are scattered throughout the genome^5^. The most common classes of RNA transposable elements in the human genome include long terminal repeat (LTR), long interspersed nuclear element (LINE), short interspersed nuclear element (SINE), and Retroposon (LINE, SINE, and Retroposon belong to non-LTR) that use RNA as their intermediate for the transposition^3^. DNA transposable elements in the human genome mainly consist of DNA transposons and RC (rolling circles or helitrons)^3^. Instead of using RNA intermediate for their transposition, DNA transposons mostly use the “cut-and-paste” mechanism for their propagation^6^, with the RC using the rolling-circle intermediates for their transposition^7^.

LINE-1 (a family of LINE class) and Alu (a family of SINE class) have been extensively studied in human genomes. LINE-1 makes up approximately 17% of the human genome, and it has been estimated that there are more than one million Alu repeats in the human genome^8^. The intact sequence for LINE-1 is about 6 kb, containing protein domains encoded by two open reading frames (ORFs), one of which (ORF2) encodes both endonuclease domain (EN) and reverse transcriptase domain (RT) that are important for the transposition. Alu is about 300 bp without the protein-coding ability, and it essentially relies on the activity of ORF2 protein encoded by LINE-1 for its transposition. In addition to Alu, SVA (SINE-VNTR(variable number of tandem repeats)_Alu, a family of retroposon class) also requires the LINE-1 for the transposition^9^. REs are generally believed to be beneficial for the species, as they can help maintain the integrity of centromeres and telomeres. They have also been extensively domesticated in the genome to benefit genome evolution^10^. In particular, most TEs in the human genome were inserted millions of years ago and have accumulated mutations and thus become defective^8^. However, it has been estimated that about 80-100 LINE-1 elements are still fully functional and capable of transposition^11^. Therefore, it is deleterious at the individual level if their activities are not adequately regulated^10^. Given the critical role REs play in chromosome integrity and genome stability, the host has evolved several mechanisms to ensure the proper regulation of REs. In humans, epigenetic silencing, including DNA methylation and histone modification, is extensively studied^6^. Biochemically, DNA methylation involves adding methyl (-CH3) group covalently to the five positions of cytosine moiety, mainly within CpG dinucleotides, which often exist in clusters called CpG islands. Enzymes, including methyltransferases (DNMTs), are responsible for DNA methylation^12^. Methyl-CpG binding domain proteins (MBD) can associate with methylated DNA, which can induce histone protein deacetylation by recruiting histone deacetylases. The interplay between DNA methylation and histone acetylation is critical for regulating chromatin conformation, which is essential for regulating gene expressions, including RE expression^12,13^. Besides DNA methylation and histone modifications, piwi proteins and their associated piwi RNA complexes have also been extensively studied for their role in the silencing of REs in the germline via either transcriptional gene regulation (*e.g.*, DNA methylation) or posttranscriptional gene regulation (*e.g.*, bind to and degrade TE transcripts in the cytosol)^11^.

### Dysregulation of REs in cancer genomes

Compared with normal cells, cancer cells typically present an aberrant epigenetic landscape where hypermethylation of promoter regions in the tumor suppressor genes is coupled with extensive hypomethylation in the intergenic regions^14^. CpG islands (regions with a high frequency of CpG sites) of gene promoters are a relatively small part of the genome compared with the vast majority of CpG dinucleotides found in the regions of REs^12^. For example, by analyzing Illumina 450K methylation array data of 10 different types of common cancer samples (including LIHC, HNSC, BLCA, LUSC, COAD, BRCA, KIRC, PRAD, LUAD, and KIRP) compared with the corresponding matched normal samples in the TCGA (The Cancer Genome Atlas) dataset, a recent study found that out of 10 tumor types, 5 of them (LIHC, HNSC, BLCA, LUSC, COAD) showed more hypo-methylated CpGs than hyper-methylated CpGs among differentially methylated CpGs. Importantly, when restricting the analysis of CpGs within TEs, 9 out of 10 analyzed cancer types (except KIRP) showed a significantly higher proportion of hypo-methylated CpGs than hyper-methylated CpGs^3^. This result indicates that the hypo-methylated TEs are the potential driving force for the extensively observed genome-wide hypomethylation in the cancer genome^12^. Furthermore, a negative relationship was observed between the intergenic TE expression and TE methylation (DNA methylation probes within +/-500 bp region around most 5’ sites of intergenic TE annotated in RepeatMasker^2^) at the subfamily level^3^. In terms of temporal activities of TEs during tumor progression, by modeling the progression of tumorigenesis via a series of cell transformations in fibroblast cells, a recent study showed that TE expressions at the subfamily level are significantly increased as cells progress through transformation (*i.e.*, increasing number of TE subfamilies are up-regulated from early passage to immortalized stage to early transformation)^15^. Notably, it was found that genome hypomethylation occurs at an early stage of transformation, and similar to findings in human cancer studies, this hypomethylation is more pronounced in TE regions^15^. Furthermore, it is observed that the methylation of these TEs remains dynamic (*e.g.*, a given subfamily of TEs can change their methylation level) during the transformation^15^. Besides TEs, studies^16,17^ also revealed the dysregulation of satellites in different types of cancers. For example, a recent study found that in bladder cancers, Sat-a (satellite-alpha) and NBL-2 (microsatellite) were hypo-methylated while D4Z4 (macrosatellite) was hyper-methylated compared with normal control; on the other hand, in leukemia, DNA methylation was increased in NBL-2 and D4Z4^16^. Finally, our previous study on osteosarcoma also found significantly higher expression levels of different satellites in osteosarcoma tumor samples compared with normal controls^17^. It is therefore clear that REs are dynamically dysregulated both epigenetically and transcriptionally in different cancer types as the tumor progresses.

Despite these advancements, dysregulation of REs in most cancer studies is restricted to the subfamily level (*i.e.*, aggregated measurement) analysis, which may limit the effectiveness of REs as biomarkers in cancer research. Recently, increasing amounts of attention^15,18,19^ have been shifted to the characterization of REs in the cancer genome at the locus-specific level. However, most studies only focused on a few types of REs (*e.g.*, HERV: human endogenous retrovirus, a member of LTR) or a few types of cancers. For instance, a recent study focusing on the locus-specific HERV in head and neck cancer patients showed that different clusters of patients based on HERV expression in the tumor-adjacent normal tissues had significantly different survival probabilities^18^.

To explore the dysregulation of locus-specific REs in cancer, we identified differentially expressed REs between tumor and the matched normal samples in 12 TCGA cancer types (including BLCA, BRCA, COAD, ESCA, HNSC, KIRC, KIRP, LIHC, LUAD, PRAD, THCA, and UCEC)^20^. We found that genic REs were co-regulated with their corresponding transcripts, defined as having overlaps in chromosomal coordinates. We identified the uniquely and recurrently dysregulated REs as well as the biological functions with their associated genes. With the recurrently dysregulated REs (REs that are dysregulated in any seven cancer types), we also identified their associated cancer genes. We identified 6 REs consistently dysregulated across all 12 cancer types: one up-regulated and five down-regulated. Notably, four of the five down-regulated REs were located within the intronic region of the TMEM252 gene, which itself was down-regulated in ten out of twelve cancer types. Our analysis of differentially expressed and methylated REs between tumor and their matched normal controls revealed a consistent negative correlation between RE methylation and expression at the locus-specific level for some cancer types.

## Material and methods

### Determine differentially expressed REs at locus-specific levels across 12 cancer types

This study analyzed twelve cancer types from the Cancer Genome Atlas (TCGA) database: BLCA, BRCA, COAD, ESCA, HNSC, KIRC, KIRP, LIHC, LUAD, PRAD, THCA, and UCEC ^20^. We selected cancer types and subjects (patients) with at least five patients per type and paired RNA-sequencing (RNA-seq) and methylation data from both tumors and matched normal samples for each patient. RNA-seq data in the bam format was downloaded using the GDC Transfer Tool Client (https://gdc.cancer.gov/access-data/gdc-data-transfer-tool). The number of subjects analyzed in each cancer type is detailed in **Table S1**.

The workflow for RE expression analysis is shown in **Fig. S1a**. Specifically, the bam files were first converted to paired-end reads in fastq format with samtools (version: 1.18)^21^. The paired-end reads were cleaned with Trim Galore (version: 0.6.4, https://github.com/FelixKrueger/TrimGalore) to remove adaptors and low-quality bases at read ends by Cutadapt (version: 4.5)^22^. The quality of clean reads was assessed with Fastqc (version: v0.11.8)^23^ before aligning to the reference human genome hg38 with STAR (version: 2.7.11a)^24^. The gene expression at the transcript isoform level and RE expression at the locus-specific level were then determined via TElocal (version 1.1.1, https://github.com/mhammell-laboratory/TElocal). Specifically, the TElocal takes alignment results generated from STAR, the gene annotation file in gtf format downloaded from UCSC Table Browser^25^, and a pre-built locus-specific RE gtf indices downloaded from https://labshare.cshl.edu/shares/mhammelllab/www-data/TElocal/prebuilt_indices/hg38_rmsk_TE.gtf.locInd.gz as inputs, and outputs the read count table for annotated transcripts and REs in the reference human genome. To determine the differentially expressed REs in each cancer type compared with matched normal samples at the locus-specific level, DESeq2^26^ was used to analyze the TElocal results with the *design = ∼ type (*tumor/normal) *+ patient_ID* to account for any individual specific effects (*i.e.*, potential confounding effects from the different characteristics of the individual patient). The normal samples were used as the reference for differential expression analysis. Specifically, count tables generated by TElocal for each cancer type were combined into a single table, which was then analyzed using the DESeq function from the DESeq2 package. REs with a |log2FoldChange| >= 1 and the adjusted p-values <= 0.05 were considered as the differentially expressed REs comparing tumors with matched normal samples. In the downstream analysis, we then focused on the locus-specific REs that belong to one of the following seven classes: DNA, LINE, LTR, RC, Retroposon, SINE, and Satellite in terms of the RE annotations (https://labshare.cshl.edu/shares/mhammelllab/www-data/TElocal/annotation_tables/hg38_rmsk_TE.gtf.locInd.locations.gz). Notably, REs with highly similar sequences, which can lead to ambiguous mapping with short reads, were excluded from further analysis. We visualized RE expression changes in each cancer type with the R package EnhancedVolcano (https://github.com/kevinblighe/EnhancedVolcano).

To understand how RE dysregulation varies across different genome regions, we categorized differentially expressed REs based on their location. First, we downloaded annotations for various genic features (5’UTRs, coding regions, introns, and 3’UTRs) in hg38 bed format from the UCSC Table Browser^25^. These were then merged using BEDTools’ mergeBed module^27^. The remaining regions of the hg38 genome were designated as intergenic using BEDTools’ complementBed module. Finally, the intersect module assigned each differentially expressed RE to a genic or intergenic category. We categorized differentially expressed REs as either genic (within genes) or intergenic (between genes) to understand how their dysregulation varies across the genome. For genic REs, we investigated the transcriptional regulation of their corresponding genes. We first identified uniquely up- and down-regulated REs specific to each cancer type within genic regions. Next, we explored the biological functions associated with the genes linked to these uniquely dysregulated REs. We used the Python package mygene (https://github.com/biothings/mygene.py) to retrieve the genes corresponding to transcripts associated with these REs, followed by functional enrichment analysis using g:Profiler^28^. For intergenic REs, we employed t-distributed Stochastic Neighbor Embedding (t-SNE) from scikit-learn^29^ to visualize distinct sample types. As a non-linear dimensionality reduction technique, t-SNE preserves the relationships between samples in high-dimensional space by maintaining these relationships in a lower-dimensional map. This allows us to visualize potential clusters or groupings within the data based on intergenic RE expression patterns.

To identify REs commonly dysregulated across multiple cancer types, we analyzed the union of up-or down-regulated REs in various combinations of cancer types, ranging from two (*i.e.*, the union of up- or down-regulated REs in any two cancer types) to twelve (*i.e.*, the union of up- or down-regulated REs in all 12 cancer types). We focused on REs that were dysregulated in at least seven cancer types (more than half of the total analyzed), refereed them as recurrently up- or down-regulated REs, and visualized them using the Python package tagore^30^. To assess the biological significance of these recurrently dysregulated REs, we analyzed the genes associated with these genic REs using g:Profiler^28^. We also intersected these genes with a curated list of 2,682 well-established cancer-related genes from COSMIC Cancer Gene Census^31^, TSGene^32^, IntOgen^33^, oncogene database^34^, and OncoKB Cancer Gene List^35^. The expression changes (log2Fold) of the relevant transcripts between the tumor and matched normal samples were visualized using the R package pheatmap^36^. Finally, we employed IGV (Integrative Genomics Viewer)^37^ to visually validate the recurrently dysregulated REs across all 12 cancer types.

### Determine differentially methylated REs at locus-specific levels across 12 cancer types

Illumina 450K methylation array offers single-base resolution methylation data for over 450,000 CpG sites across the human genome^38^. While covering 96% of CpG islands and previously identified differentially methylated regions in cancer (https://www.illumina.com/content/dam/illumina-marketing/documents/products/datasheets/datasheet_humanmethylation450.pdf), it only targets about 1.5% of all CpG sites^39^. Specifically, the 50 bp probes employed by this array can cover 99% of RefSeq genes, including their gene bodies and promoter regions. Similar to RNA-seq data, methylation data (SeSAMe Methylation Beta Values from Methylation Array Harmonization Workflow) expressed in beta values for each subject were downloaded using the GDC Transfer Tool Client (https://gdc.cancer.gov/access-data/gdc-data-transfer-tool). The workflow for RE methylation analysis is shown in **Fig. S1b**. Specifically, the hg19 annotation associated with each probe was obtained via the R annotation package IlluminHumanMethylation450kann.ilmn12.hg19 (http://www.bioconductor.org/packages/ <u>IlluminaHumanMethylation450kanno.ilmn12.hg19/</u>)^40^ and the conversion of the coordinates from hg19 to hg38 was conducted using pyliftover (https://pypi.org/project/pyliftover/), where only the probes that can be converted to hg38 were kept for further analysis. Focusing on the transcription start site (TSS) known to be associated with transcriptional repression in cancer^41^, we defined a 1kb region flanking the 5’ end (TSS +/- 500 bp) of each RE in terms of the previous study^42^. We used pybedtools^27,43^ to identify probes intersecting these 1kb regions, ensuring they mapped uniquely to each RE locus. Probes that can be mapped to multiple RE loci were removed to reduce the ambiguity. Average beta values were calculated for RE loci mapped by multiple probes to represent their methylation levels.

Similar to RE expression analysis, we focused on REs belonging to the seven RE classes. Differentially methylated REs were identified using the limma R package^44^, with the *design = ∼ type (*tumor/normal) *+ patient_ID* to account for individual effects^42^. Beta values were converted to M values (M value = log2 (Beta value / (1 – Beta value))) for the statistical tests given its approximate normal distributional properties. Differentially methylated REs were identified based on the adjusted p-value <= 0.05 and an absolute difference in beta values between tumor and matched normal samples >= 0.1, as in the previous study^42^. Due to the limited coverage of Illumina 450K methylation array on the human genome, methylation data was primarily used for association analysis with RE expression data, as described below.

### Determine the association between RE methylation and expression changes at the locus-specific level

While a previous study identified a negative association between the intergenic TE expression and the methylation level at the subfamily level^3^ we aimed to examine this relationship at the locus-specific level. In this study, however, only 66,859 of the 4,467,488 locus-specific REs for expression analysis have corresponding methylation probes (107,634 probes). The limited overlap between differentially expressed and methylated REs necessitates separate analyses. For each cancer type, we compared the expression changes (log2 fold changes) between hypo- and hyper-methylated REs. Additionally, we compared the average methylation changes (tumor vs normal) based on M values for up- and down-regulated REs. Finally, the Pearson correlation coefficient between the methylation changes (*i.e.*, Tumor - Normal) based on M values and expression changes based on log2 fold changes of normalized expressions (*i.e.*, log2 (Tumor/Normal)) for each of these REs was determined based on all subjects in a given cancer type.

## Statistical analysis

To assess the statistical significance of differences in expression and methylation changes between tumor and matched normal samples, we employed the non-parametric Wilcoxon rank-sum test implemented in the R package ggpubr^45^. A p-value < 0.05 was considered statistically significant. For enrichment analysis of gene sets, we used the hypergeometric test implemented in g:Profiler^28^. Only annotated genes were included in the statistical domain scope. Multiple testing correction was performed using the default g:SCS method, and adjusted p-values < 0.05 were considered significant. Pearson correlation coefficients were calculated using the pearsonr function implemented in SciPy^46^ for the correlation analysis between RE methylation and expression changes.

## Results

### Differentially expressed REs between tumor and matched normal samples at the locus-specific level

Our analysis focused on locus-specific REs in the human genome, as annotated by TElocal for hg38. There are 4,505,469 locus-specific REs, mostly belonging to SINE, LINE, LTR, and DNA classes (**Fig. S2a** and **2b**). Sequence lengths varied across 7 RE classes: RC, SINE, DNA, and Retroposons ranged from 10 to 1,000 bps, while LINEs spanned 10 to 10,000 bps, and Satellite and LTR elements ranged from 10 to 100,000 bps (**Fig. S3a**). The number of REs generally corresponded to chromosome size across the 22 autosomes (**Fig. S3b**). About half of the human genome comprises various REs (**Fig. S3c**). Furthermore, most REs associated with protein-coding and non-protein-coding genes (based on RefSeq annotation) reside in intronic or intergenic regions (**Fig. S3d**). Finally, we excluded 37,981 REs with identical genomics sequences to reduce ambiguity in expression analysis, resulting in 4,467,488 REs for further analysis.

Based on these 4,467,488 locus-specific REs, we analyzed RE dysregulation patterns in 12 cancer types, identifying distinct expression changes for each. Concretely, the differentially expressed REs between tumor and matched normal samples in each cancer type were determined separately. As shown in **Fig. S4** and **Table S2**, KIRC exhibited the highest number of up-regulated REs (40,125), followed by BRCA (16,586). Conversely, BRCA had the most down-regulated REs (45,586), followed by THCA (31,099). Interestingly, UCEC and HNSC showed the fewest up- and down-regulated REs, respectively. A detailed breakdown of these differentially expressed REs in either genic or intergenic regions is shown in **Fig. 1a**. Clearly, most of these differentially expressed REs fall into the largest 4 RE classes (*i.e.*, SINE, LINE, LTR, and DNA) in the genic regions. Interestingly, among all RE classes, HNSC has a relatively higher number of Retroposon being up-regulated in tumor samples compared to the matched normal controls (*i.e.*, 720 genic RE up-regulations + 601 intergenic RE up-regulations vs 5 genic RE down-regulations + 4 intergenic RE down-regulations, see **Table S2** for detailed information in the context of other cancer types).

**Fig. 1:**
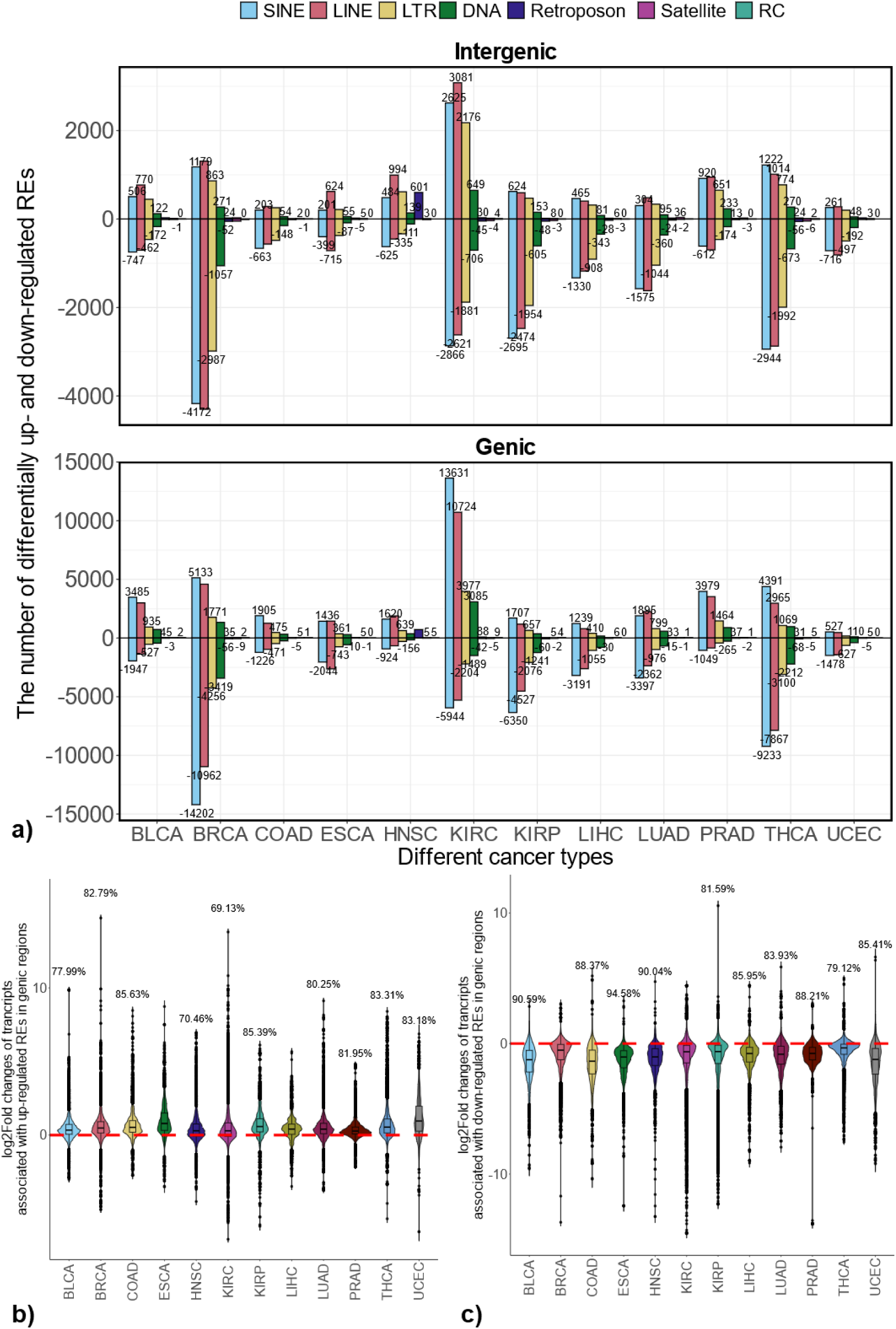
Differentially expressed locus-specific REs across 12 cancer types and co-regulation of transcripts with corresponding genic REs. **a):** number of up- or down-regulated REs stratified by their genomic contexts; **b):** expression changes of transcripts associated with corresponding up-regulated genic REs. **c):** expression changes of transcripts associated with corresponding down-regulated genic REs.

For both up- and down-regulated genic REs, we observed a strong co-regulation with their corresponding transcripts across all 12 cancer types. As shown in **Fig. 1b**, for up-regulated genic REs, the large proportions of the log2Fold changes for the associated transcripts show positive values across all 12 cancer types (indicating up-regulation). However, these proportions varied slightly among different cancer types. A similar co-regulation between RE and associated transcripts is also consistently observed for the down-regulated genic REs, with predominantly negative values in log2 fold change, as shown in **Fig. 1c**.

### Uniquely differentially expressed REs in each cancer type at the locus-specific level

To identify REs uniquely differentially expressed in each cancer type, we analyzed the differentially expressed REs identified in each of the 12 cancer types (see the details in **Table S3-S14** for uniquely dysregulated REs in BLCA, BRCA, COAD, ESCA, HNSC, KIRC, KIRP, LIHC, LUAD, PRAD, THCA, and UCEC respectively). This analysis revealed that each cancer type exhibits a distinct pattern of RE dysregulation, with unique sets of REs being up- or down-regulated. **Figures 2a** and **2b** summarize the number of uniquely up- and down-regulated REs stratified by their genomic context (genic vs. intergenic) in each cancer type. As expected, KIRC, which had the highest overall number of up-regulated REs, also exhibited the most uniquely up-regulated REs, primarily in genic regions. Similarly, BRCA and THCA demonstrated the highest number of uniquely down-regulated REs, predominantly within genic regions.

**Fig. 2:**
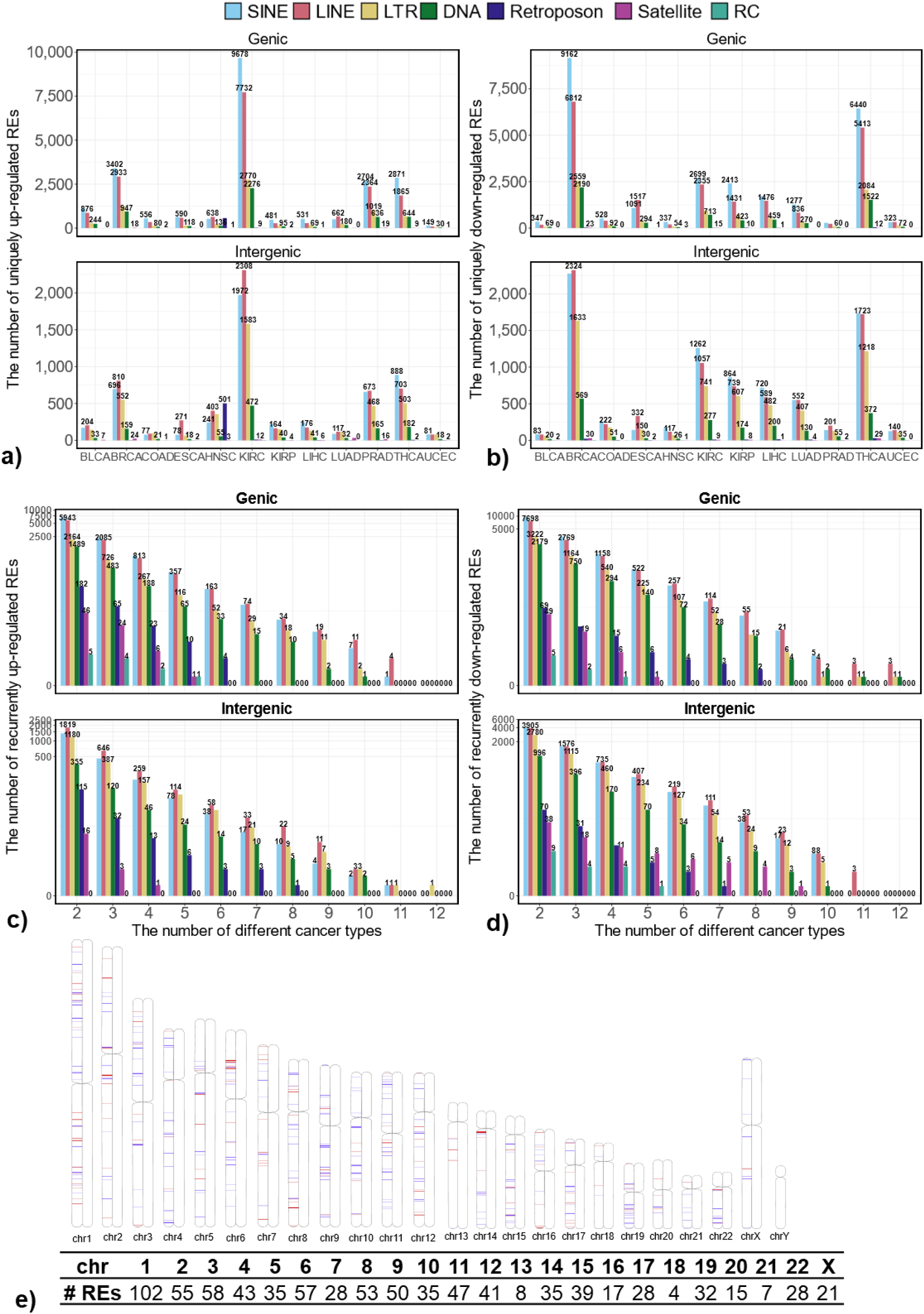
Uniquely and recurrently dysregulated REs across 12 cancer types. **a):** number of uniquely up-regulated REs in a given cancer type stratified by their genomic context; **b):** number of uniquely down-regulated REs in a given cancer type stratified by their genomic context; **c):** number of recurrently up-regulated REs among any number of 12 cancer types stratified by their genomic context; **d):** number of recurrently down-regulated REs among any number of 12 cancer types stratified by their genomic context; **e):** Genomic locations of recurrently up- (red lines) and down-regulated (blue lines) REs in any seven cancer types (defined as the recurrently dysregulated REs in this study) and their abundance in each human chromosome.

To understand the biological relevance of uniquely differentially expressed REs, we performed gene set enrichment analysis for genes associated with these elements in genic regions. Briefly, we retrieved genes linked to uniquely dysregulated genic REs using the mygene package (https://github.com/biothings/mygene.py). The relationship between the number of REs, associated transcripts, and corresponding genes is summarized in **Table S15**. We then used g:Profiler^28^ to analyze the functional terms associated with these genes, including Gene Ontology (GO) terms of GO: MF, GO: BP, and GO: CC and terms related to biological pathways in KEGG, Reactome, and WikiPathways. **Figures 3a** and **3b** depict the top five enriched functional terms for uniquely up- and down-regulated REs across the 12 cancer types. All terms were statistically significant (adjusted p-values < 0.05), based on the g_SCS correction method. The rich factor is calculated as the intersection size (*i.e.*, the number of genes in the input query annotated to the corresponding term) divided by the term size (*i.e.*, the number of genes annotated to the term in hg38 genome annotation), multiplied by 100 (%).

**Fig. 3:**
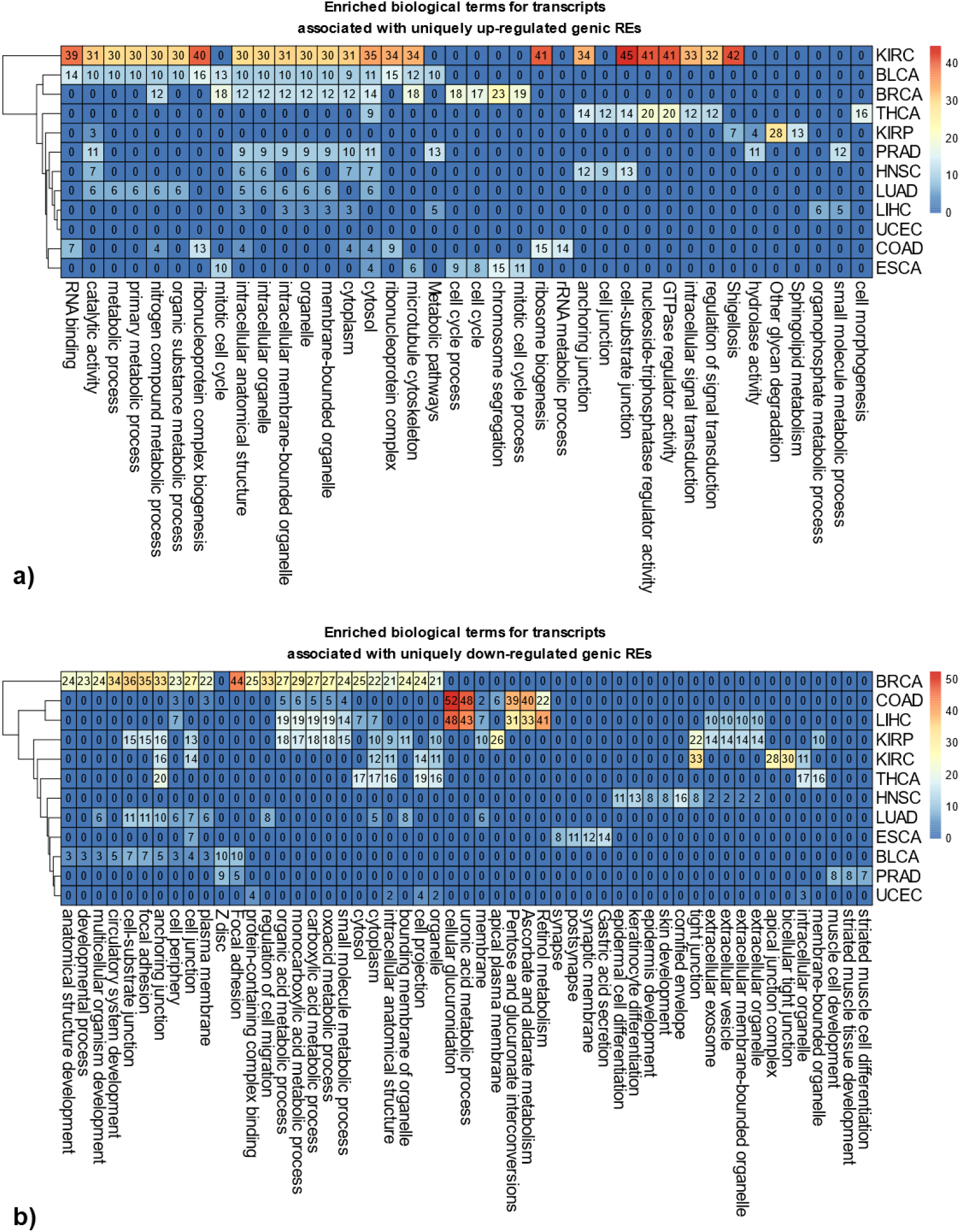
Enriched biological functions associated with uniquely dysregulated genic REs. **a):** top 5 enriched functional terms, including gene ontology (GO) terms (including GP: MF, GO: BP, and GO: CC) and terms associated with biological pathways (including KEGG, Reactome, and WikiPathways) associated with uniquely up-regulated genic REs. **b):** top 5 enriched functional terms associated with uniquely down-regulated genic REs. (note: numbers in the heatmap indicate the rich factors calculated as ((number of genes in the input query that are annotated to the corresponding term) / (number of genes that are annotated to the term)) X 100%).

In terms of uniquely up-regulated genic REs, as shown in **Fig. 3a**, among a total of 38 (*i.e.*, union of top 5 enriched terms from all 12 cancers) functional terms, 24 of them are significantly enriched in KIRC, followed by 18 terms in BLCA and 14 terms in BRCA. Notably, UCEC lacked enriched terms. Namely, genes associated with uniquely up-regulated genic REs in UCEC do not contain any enriched terms from GO and Pathway databases. Interestingly, BRCA and ESCA are enriched in the terms related to the cell cycle process, whereas KIRC, BLCA, BRCA, PRAD, HNSC, LUAD, and LIHC showed enrichment in intracellular structures, with KIRC and THCA also displaying enriched terms in signal transduction pathways. Among 53 functional terms associated with uniquely down-regulated genic REs, BRCA had the most enriched terms (24), followed by KIRP (21). ESCA, PRAD, and UCEC each had 5 enriched terms (**Fig. 3b**). BRCA, KIRP, LUAD, BLCA, and, to a lesser extent, KIRC, THCA, and ESCA all displayed enrichment in cell junctions. COAD and LIHC showed enrichment in distinct pathways, including cellular glucuronidation, uronic acid metabolic process, pentose and glucuronate interconversion, ascorbate, aldarate, and retinol metabolism.

REs that are uniquely differentially expressed in each cancer may represent the unique characteristics of that cancer. To assess the potential of uniquely differentially expressed REs as cancer-type representations, we focused on intergenic REs to minimize confounding effects from associated transcripts. We used t-SNE^29^ to visualize the normalized expression of these REs, including uniquely up-regulated, down-regulated, and combined sets. Briefly, the normalized expression of uniquely up- or down-regulated intergenic REs, as well as the combination of them, were used as the feature inputs to t-SNE to cluster different sample types (including different normal samples, different tumor samples, different normal and tumor samples) as shown in **Fig. 4**. Specifically, a total of 17,541 uniquely up-regulated intergenic REs, 24,351 uniquely down-regulated intergenic REs, as well as 38,899 uniquely dysregulated intergenic REs (*i.e.*, the union of up-regulated and down-regulated REs, there are 2,993 REs that are overlapped between uniquely up-regulated and down-regulated REs because some of the uniquely up-regulated REs in one cancer can be down-regulated in other cancers) were used as the input for t-SNE visualizations (see **Fig. 4a-b** and **Fig. S5a-f**). Clearly, **Fig. 4** demonstrates the clustering of different sample types (normal, tumor, and combinations) based on these REs. Notably, uniquely up-regulated intergenic REs effectively differentiate sample types of different tissue origin for both normal and tumor samples, as shown in **Fig. 4a**. Interestingly, the tumor and their matched normal samples are close together. Still, they are distinct from other sample types in this two-dimensional latent space when these uniquely up-regulated intergenic REs are used to cluster different normal and tumor samples, as shown in **Fig. 4b**. Similar trends were observed with uniquely down-regulated and combined intergenic REs (**Fig. S5c-f**).

**Fig. 4:**
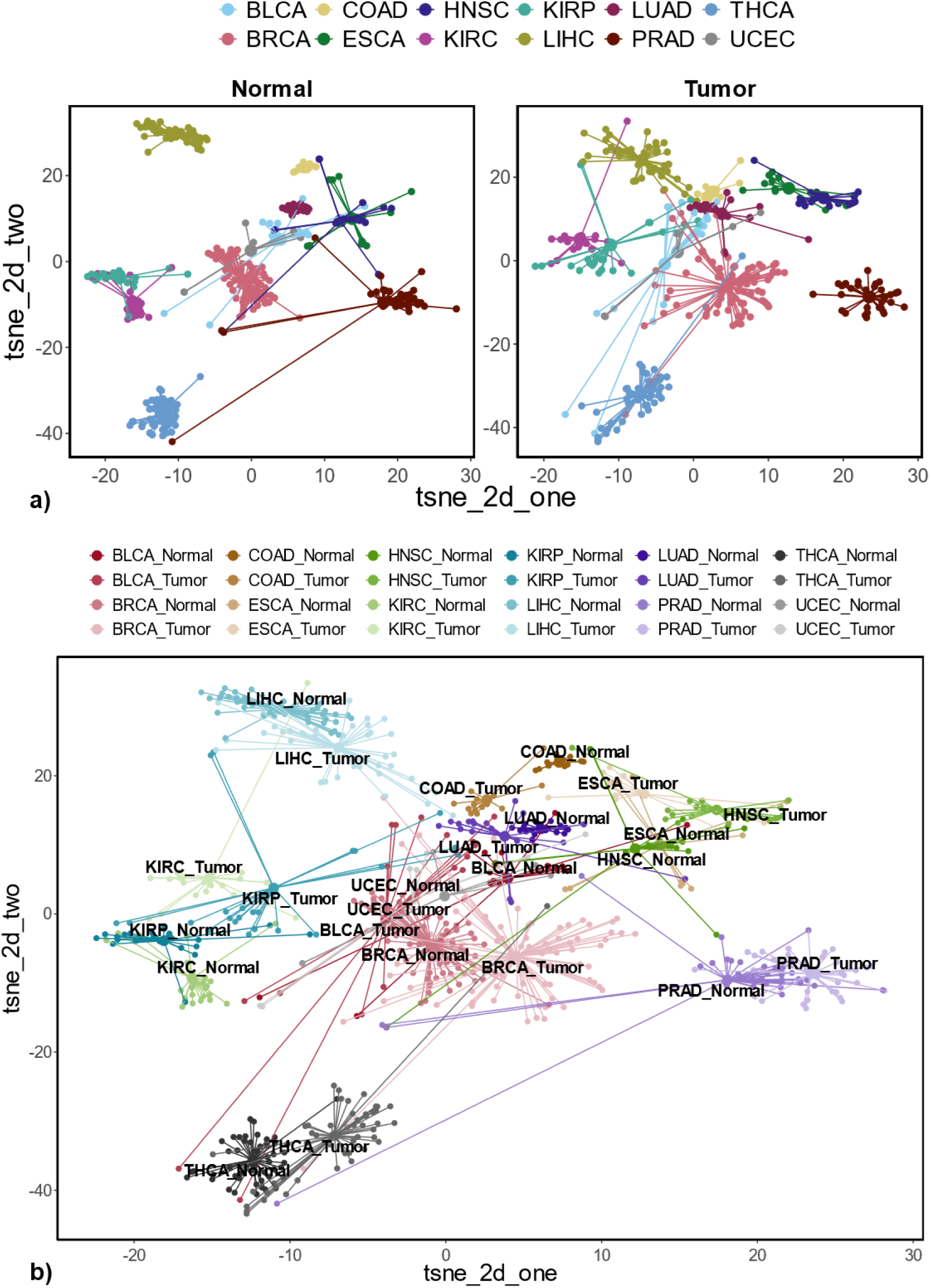
Sample clustering with uniquely up-regulated intergenic REs identified across 12 cancer types. (tsne_2d_one: first dimension, tsne_2d_two: second dimension)**. a):** t-SNE plots based on uniquely up-regulated intergenic REs for Normal and Tumor sample clustering separately; **b):** t-SNE plot based on uniquely up-regulated intergenic REs for different sample type clustering (tumor with matched normal samples);

### Recurrently differentially expressed REs in multiple cancer types at the locus-specific level

To identify REs commonly dysregulated across 12 cancer types, we analyzed the overlap of up- and down-regulated REs in various combinations. We focused on those appearing in at least seven cancer types (more than half of the 12 cancer types), which we termed “recurrently differentially expressed REs” (see the details in **Table S16**). As shown in **Fig. 2c**, we identified a total of 272 recurrently up-regulated REs, including 188 in genic regions (15 DNA, 74 LINE, 29 LTR, 70 SINE) and 84 in intergenic regions (10 DNA, 33 LINE, 21 LTR, 17 SINE, 3 Retroposon). Similarly, we identified 566 recurrently down-regulated REs (**Fig. 2d**), with a similar distribution between genic and intergenic regions, namely, 295 in genic regions (consisting of 28 DNA, 114 LINE, 52 LTR, 98 SINE, 3 Retroposon REs) and 271 in intergenic regions (composed of 14 DNA, 111 LINE, 54 LTR, 5 Satellite, 86 SINE, 1 Retroposon). Clearly, most of these recurrently dysregulated REs belong to LINE, SINE, LTR, and DNA classes. **Figure 2e** illustrates the genomic locations of these recurrently up- and down-regulated REs within each chromosome. Notably, chromosome 1, the largest human chromosome, contains the highest number of recurrently dysregulated REs (102 out of 838 or 12.17%).

Moreover, we found that among the transcripts associated with recurrently up-regulated genic REs, a high percentage exhibited increased expression in tumor samples compared to matched normal controls across all 12 cancer types (**Fig. 5a**). Specifically, 188 recurrently up-regulated genic REs are associated with 588 transcripts. Among these transcripts, 90.14% in BLCA show a positive log2 fold change in expression, while BRCA has the lowest percentage at 79.08%. The remaining cancer types fall between these two values. Similarly, transcripts associated with recurrently down-regulated REs displayed decreased expression in tumors. We detected 295 recurrently down-regulated genic REs that correspond to 863 transcripts, most of which are down-regulated, as shown in **Fig. 5b**. Among these transcripts, 90.96% in ESCA exhibit a negative log2 fold change when comparing tumor and matched normal samples, while PRAD has the lowest percentage at 71.15%. The remaining cancer types fall between these two.

**Fig. 5:**
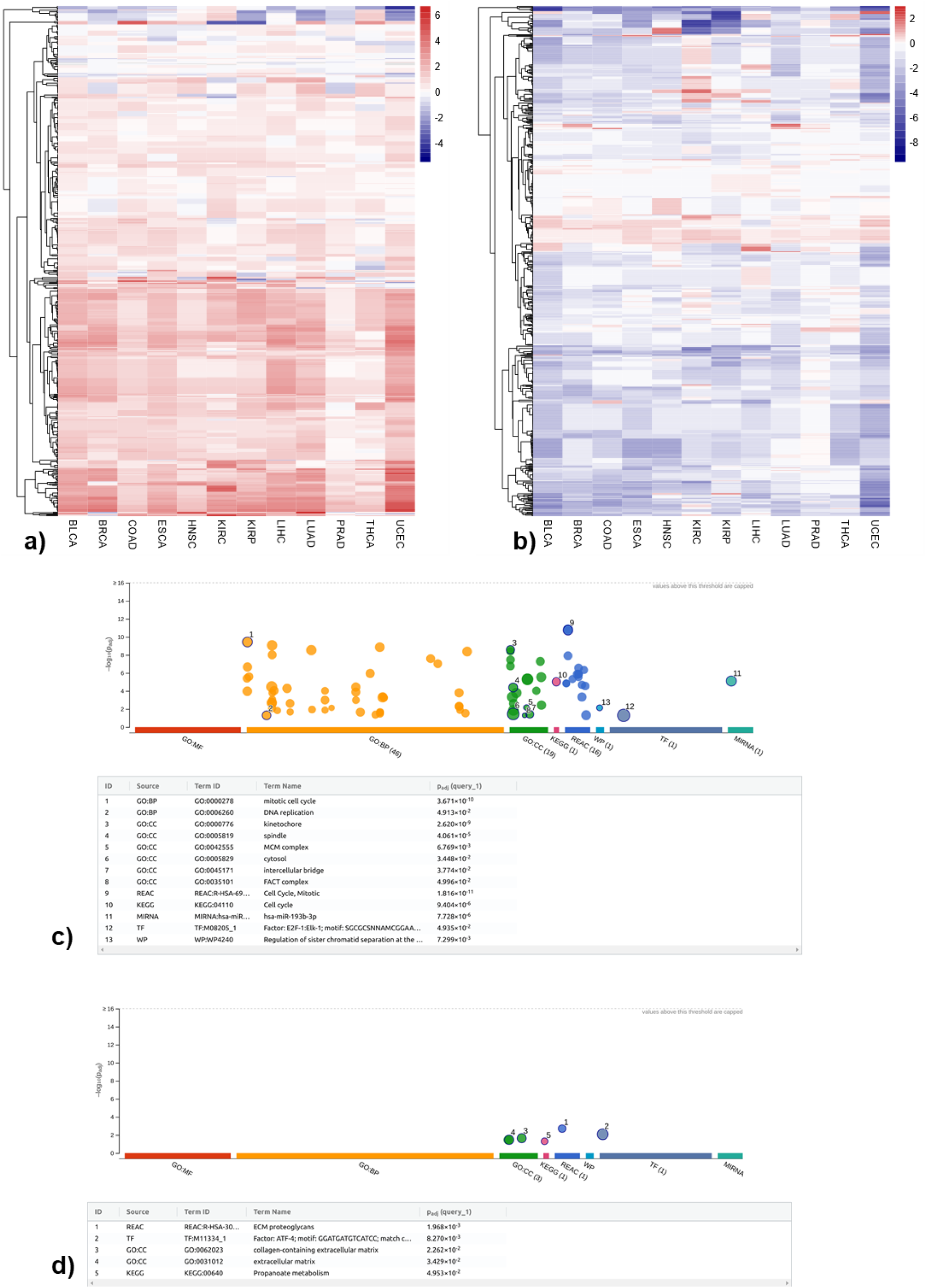
Regulation and functions of transcripts associated with recurrently dysregulated genic REs. **a):** log2 expression changes between tumor and matched normal samples for transcripts corresponding to recurrently up-regulated genic REs; **b):** log2 expression changes between tumor and matched normal samples for transcripts corresponding to recurrently down-regulated genic REs; **c):** enriched biological functions associated with the genes in a; **d):** enriched biological functions associated with the genes in b.

Using the mygene package, we retrieved genes for transcripts associated with recurrently up- and down-regulated genic REs. Among the 588 transcripts associated with recurrently up-regulated genic REs, 450 correspond to 121 genes, while the remaining 138 transcripts lack corresponding reference genes. Similarly, among the 863 transcripts associated with recurrently down-regulated genic REs, 658 correspond to 158 genes, with the remaining 205 transcripts lacking reference genes. These two gene sets were submitted to g:Profiler^28^ for functional analysis. Interestingly, the functional analysis revealed distinct patterns for up-vs. down-regulated REs. Genes associated with recurrently up-regulated REs were enriched for functions in the cell cycle and DNA replication (**Fig. 5c**), processes crucial for cancer cell proliferation. Conversely, genes associated with recurrently down-regulated REs displayed enrichment for extracellular matrix (ECM) and ECM proteoglycans (**Fig. 5d**), suggesting potential tumor microenvironment remodeling.

To further investigate the potential importance of these genes in tumor development, we compared these two sets of genes to a curated list of 2,682 well-established cancer genes, as mentioned in the method section. We identified 12 cancer genes associated with recurrently up-regulated REs (CDK1, ESCO2, GAS5, GTSE1, H2BC12, KIFC1, MMS22L, NF1, NIT2, PVT1, STAT1, UBE2C) and 19 cancer genes associated with recurrently down-regulated REs (ADAMTS9-AS2, ATF3, CASP8, DCN, DUSP1, ECT2L, EMP1, FANCC, JDP2, KLF6, NDRG2, NR4A1, RHOBTB2, SOCS2, SPARCL1, STARD13, SYNPO2, TAGLN, TIMP3). The expression changes of the cancer-related transcripts associating with recurrently up- and down-regulated genic REs between tumor and matched normal controls are shown in **Fig. S6a** and **S6b**, respectively. While most genes associated with recurrently dysregulated REs are transcriptionally regulated in a similar manner (up-regulated REs associated with up-regulated genes, and vice versa, see **Fig. 1b and 1c**), some genes, such as CASP8, FANCC, ECT2L, and SPARCL1, exhibit opposing expression patterns. For example, in the case of the CASP8 annotated with nine transcripts, 105 REs were found in its genic regions. Although some of these REs (*e.g.*, SVA_E_dup106, SVA_E_dup107) are recurrently down-regulated in most of the 12 cancer types, the vast majority of these 105 REs are up-regulated, as shown in **Fig. S6c**. A similar scenario also holds for the FANCC, as shown in **Fig. S6d**. On the other hand, for gene ECT2L and especially SPARCL1, their up-regulation shown in **Fig. S6b** is indeed consistent with the up-regulation of the corresponding genic REs shown in **Fig. S6e** and **S6f,** respectively. This suggests complex regulatory mechanisms might be involved in RE expression and regulations.

Beyond the recurrently dysregulated REs identified in at least seven cancer types, we also identified a subset of REs consistently dysregulated in all 12 cancer types. As shown in **Fig. 2c**, one intergenic RE (MLT1D_dup1540), which belongs to the MLT1D subfamily, ERVL-MaLR family, and LTR class, is consistently up-regulated in all 12 cancers comparing tumors with matched normal controls. Furthermore, five genic REs (*i.e.*, MamTip2_dup3664, L1MB2_dup4373, L1ME3Cz_dup10031, MamRTE1_dup5302, and LTR82A_dup581) are consistently down-regulated in all 12 cancer types, as shown in **Fig. 2d**. We validated these findings using IGV, comparing read coverage between tumor and matched normal samples. An example of read coverage corresponding to MLT1D_dup1540 between 5 randomly selected tumors and matched normal samples from BLCA (a total of 17 paired tumor-normal samples in BLCA) is shown in **Fig. S7a**. The comparison of normalized expression levels across all 12 cancer types is shown in **Fig. S7b**, which is consistent with the IGV results. Interestingly, four of these five consistently down-regulated REs (except for L1ME3Cz_dup10031) are located in the same intronic region of the TMEM252 gene (with transcripts ID: NM_153237.2, consisting of two exons). For example, in **Fig. 6a**, we showed the reads coverage corresponding to these 4 REs between the same randomly selected tumor and matched normal samples from BLCA (note that no reads are mapped to the 5^th^ RE: LTR82A_dup582). The log2Fold changes of TMEM252, as well as the five associated REs (displayed in **Fig. 6a**) across 12 cancer types, is depicted in **Fig. 6b**, and the corresponding normalized expression comparison between tumor and matched normal samples in BLCA is shown in **Fig. 6c**. A similar comparison across all 12 cancer types is shown in **Fig. S8**. Among these five consistently down-regulated genic REs, L1ME3Cz_dup10031 is located in the intronic region of gene GCOM1 and MYZAP. The IGV genomic context and the normalized expression comparison for L1ME3Cz_dup10031 between tumor and matched normal samples across all 12 cancer types are shown in **Fig. S9**.

**Fig. 6:**
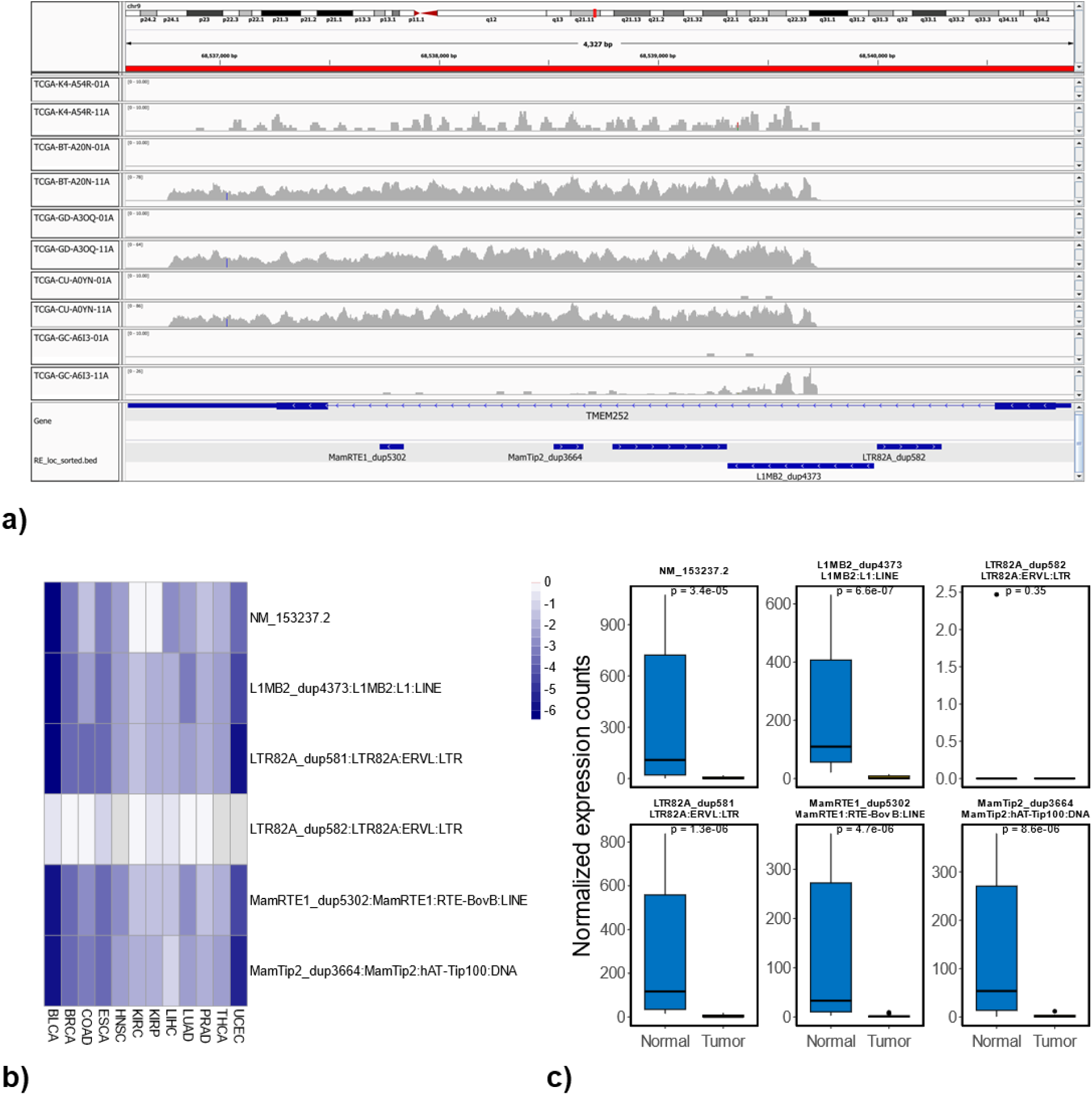
Consistent down-regulated 4 REs and their corresponding gene TMEM252 among 12 cancer types. **a):** reads coverage corresponding to these 4 REs (down-regulated in all 12 cancer types) and their associated gene: TMEM252 (with transcript ID: NM_153237.2) between the randomly selected 5 tumor and matched normal samples in BLCA (same set of samples as used in **Fig. S7a**, tumor sample ends with 01A while the matched normal sample ends with 11A); **b) :** The log2Fold changes of TMEM252 as well as the associated REs across 12 cancer types; **c) :** Normalized expression comparison between tumor and matched normal samples in BLCA for TMEM252 and associated REs.

### Differentially methylated REs between tumor and matched normal samples at the locus-specific level

To analyze RE methylation, we focused on probes that mapped uniquely to the 1kb region surrounding the transcription start site (TSS) of locus-specific REs. Among the 4,467,488 locus-specific REs available for expression analysis, 131,322 were covered by at least one probe within their 1kb TSS. After excluding probes that mapped to multiple RE loci (due to the overlapping REs in the genome), we identified 66,859 unique REs associated with a total of 107,634 probes (multiple probes can be mapped to the 1kb TSS of the same RE). These unique REs consist of 33,590 SINE, 17,348 LINE, 7,427 LTR, 7,988 DNA, 245 Retroposon, 241 Satellite, and 20 RC, as shown in **Fig. S10a**. The breakdown of these REs at the family levels is shown in **Fig. S10b**.

To identify differentially methylated REs between tumors and matched normal control for these 66,859 REs, the calculated M values for each RE were used as the input for the limma R package^44^. **Table S17** summarizes the number of hypo-methylated REs (*i.e.*, reduced methylation level in tumors) and hyper-methylated REs (*i.e.*, increased methylation level in tumors) in each cancer type. 10 of 12 cancer types (excluding KIRP and PRAD) exhibited significantly more hypo-methylated REs than hyper-methylated ones. Like differentially expressed REs, most differentially methylated REs belonged to SINE, LINE, LTR, and DNA classes.

### Relationship between RE methylation and expression at the locus-specific level

Given the limited coverage of Illumina 450K methylation array data, we focused on assessing the association between RE methylation and expression changes among the differentially methylated REs (see **Table S17)**. We observed a slight overlap between hypo-methylated REs and differentially expressed REs across all 12 cancer types (**Fig. S11a**). Similar patterns were found for hyper-methylated REs and differentially expressed REs (**Fig. S11b**). Furthermore, we also identified the differentially methylated REs that are up- or down-regulated in their expression. As shown in **Fig. S11c** and **S11d**, similar small overlaps were also observed in this case.

To examine the relationship between methylation and expression changes, we compared expression changes (log2 fold changes) of hypo- and hyper-methylated REs, assuming hypo-methylated REs are expected to have a higher expression change compared to hyper-methylated REs if DNA methylations negatively regulate the expression of REs. As shown in **Fig. 7a**, 4 cancer types (BRCA, KIRC, KIRP, PRAD) showed this expected pattern, while the remaining eight had non-significant results. Furthermore, we compared the averaged methylation changes (*i.e.*, Tumor - Normal) based on M values between up- and down-regulated REs with the assumption that averaged methylation changes should be lower for up-regulated REs compared to down-regulated ones. As shown in **Fig. 7b**, 6 cancer types (BRCA, COAD, KIRC, KIRP, LUAD, and PRAD) showed lower averaged methylation changes when these REs were up-regulated. Therefore, based on this two-complementary analysis, BRCA, KIRC, KIRP, and PRAD demonstrated a consistent negative relationship between RE methylation and expression at the locus-specific level. For recurrently dysregulated REs, we found limited overlap with methylation data due to array coverage. Unfortunately, the six consistently dysregulated REs identified in this study are not covered by the Illumina 450K methylation arrays. Therefore, we focused on those recurrently dysregulated in any seven cancer types. Among 272 recurrently up-regulated REs, only 16 were covered by the DNA methylation array; among 566 recurrently down-regulated REs, only 10 were covered. The Pearson correlation between methylation and expression changes for recurrently up-regulated REs is shown in **Fig. 7c**, while **7d** showed the results for recurrently down-regulated REs. While the correlation varied slightly between cancer types, a clear negative correlation was observed for some specific REs (*e.g.*, L1M4_dup16795 in **Fig. 7d**).

**Fig. 7:**
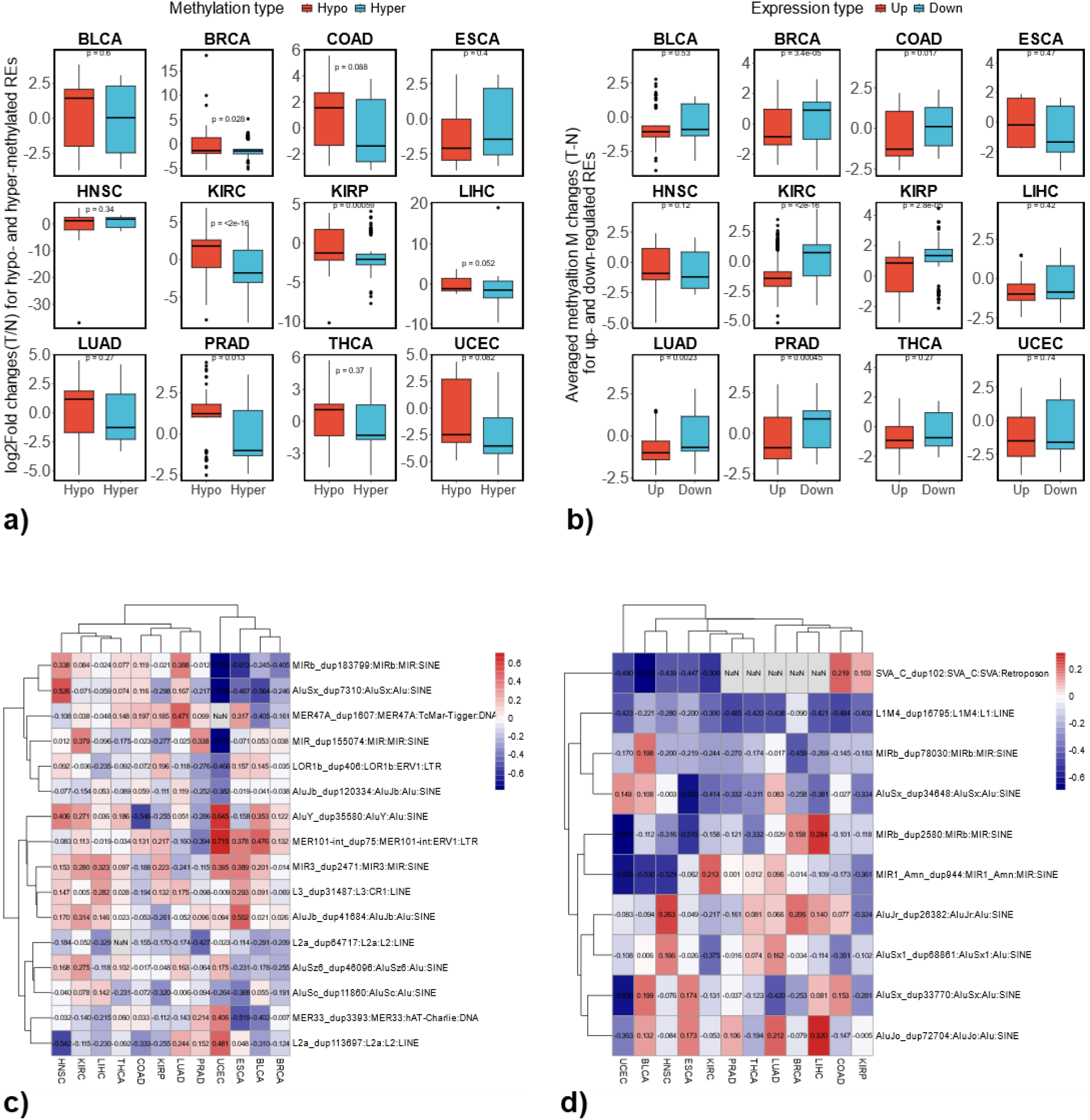
Association between changes of RE methylation and RE expression across 12 cancer types. **a):** comparison of expression changes for differentially expressed REs that are either hypo- or hyper-methylated; **b):** comparison of methylation changes for differentially methylated REs that are either up- or down-regulated; **c):** Pearson correlation coefficient between methylation changes (*i.e.*, Tumor - Normal) based on M values and expression changes based on log2 fold changes of normalized expressions (*i.e.*, log2(Tumor/Normal)) for recurrently up-regulated REs that are covered by DNA methylation array; **d):** Pearson correlation coefficient between methylation changes and expression changes for recurrently down-regulated REs that are covered by DNA methylation array;

## Discussion

Previous research has extensively explored the dysregulation of repetitive elements (REs), particularly transposable elements (TEs), in various adult and pediatric cancers^17^. Differentially expressed TEs were investigated comprehensively among 13 cancer types^42^. Although these studies primarily analyzed REs at the subfamily level, few studies investigated the dysregulation of specific REs at the more detailed, locus-specific level. For example, using Telescope^47^, a recent study compared the expression level of locus-specific HERVs between prostate, breast, and colon cancers and their matched normal controls^19^. They found that 155 HERV loci were differentially expressed in all three cancer types, and 114 were differentially expressed in the same direction. Focusing on head and neck cancer and using Telescope, expression levels of HERVs in the tumor-adjacent normal tissue were shown to help cluster patients with different survival probabilities^18^.

To better understand the dysregulation landscape of REs comprehensively at the locus-specific level in cancer genomes, we used TElocal (https://github.com/mhammell-laboratory/TElocal). Unlike Telescope, designed primarily for locus-specific HERV analysis^47^, TElocal performed better in a recent benchmarking comparison among TE RNA-Seq analysis tools^48^. Our study comprehensively investigated the dysregulation of repetitive elements (REs) at the locus-specific level across 12 cancer types, offering a more detailed picture of RE dysregulation in cancer genomes. We observed uniquely dysregulated REs in each cancer type and commonly dysregulated REs across multiple types. The dysregulation of genic REs, specifically those situated in introns, may influence the expression of corresponding genes. Notably, several genes associated with these dysregulated REs are well-known tumor suppressors or oncogenes, highlighting the potential role of REs in cancer development.

Our analysis identified distinct patterns of RE dysregulation across the 12 cancer types. After accounting for the potential confounding effects of individual subjects in our expression analysis, our results (see **Table S2**) showed that among the 12 cancer types analyzed, 5 of them (BLCA, COAD, HNSC, KIRC, PRAD) showed more up-regulated REs than down-regulated ones at the locus-specific level. In comparison, despite focusing on the intergenic TE dysregulations at the subfamily levels, the previous study using the TCGA dataset also showed the over-expression of TEs in these five cancer types^42^. On the other hand, the remaining seven cancer types (BRCA, ESCA, KIRP, LIHC, LUAD, THCA, and UCEC) showed more down-regulated REs than up-regulated ones. Among these seven cancer types, 4 of them (KIRP, LUAD, BRCA, and THCA) also showed more down-regulated intergenic TE expressions comparing tumors with matched normal samples from the same previous study^42^, where ESCA and UCEC were not analyzed. Interestingly, in the previous study^42^, LIHC displayed more up-regulated intergenic TE expressions in tumor samples compared to matched normal samples. This discrepancy is possibly due to the differences in analysis methods (*e.g.*, subfamily level versus locus-specific level), the inclusion of genic REs in our study, and the consideration of potential individual effects in our data analysis. Despite the small differences, our results are, in general, consistent with the previous findings in terms of the amount of dysregulated REs identified from different cancer types (The vast majority of REs are TE see **Fig. S2**). Compared with the previous study^42^, we also included the genic region REs in our analysis, which can help us to characterize better the biological effects of RE dysregulations in cancers by functional analysis of the associated genes. Most of these genic REs are located in the intronic regions (see **Fig. S3**), and we showed that genic REs are primarily regulated similarly to their corresponding transcripts (see **Fig. 1b-c** and **Fig. S6c-f**). This co-regulation may be because these dysregulated genic REs could be read-through transcriptions of host genes, as suggested by the previous study^42^. However, it is also possible that these REs are independent entities, regulated in a manner similar to their corresponding transcripts and genes.

To understand the unique characteristics of each cancer type, we identified REs that were exclusively up- or down-regulated in specific cancers. By analyzing the genes associated with these REs, we discovered distinct biological functions enriched in each cancer type. For example, genes linked to up-regulated REs in BRCA and ESCA were enriched in the mitotic cell cycle process, while those related to down-regulated REs in COAD and LIHC were enriched in cellular glucuronidation. This highlights how the dysregulation of genic REs, likely reflecting the dysregulation of their corresponding transcripts, can contribute to the unique features of different cancer types. In addition to the uniquely dysregulated genic REs, by projecting the uniquely dysregulated intergenic REs to a lower dimensional space with t-SNE, we demonstrated their usefulness in clustering different sample types (cancer types). Notably, these low-dimensional representations captured both the critical difference between different sample types (*e.g.*, clustering different normal sample types as well as different tumor types) as well as the difference for different tissues (*i.e.*, tumor and matched normal samples tend to cluster together, see **Fig. 4b**, **Fig. S5d,** and **S5f**). The importance of the information contained in dysregulated REs has also been recently shown to lead to improved cancer classifications for liver and esophagus cancer when differentially expressed REs measured at the subfamily level from the blood plasma of different cancer patients are included as features in a logistic regression model^49^. Compared with aggregated measurement at subfamily levels, measurement of locus-specific REs can provide richer and more informative insights into cancer studies. For example, the uniquely dysregulated intergenic REs identified in this study across different cancer types could help us better characterize each cancer type.

In addition to the uniqueness of each cancer type, we also explored their commonality by determining the commonly dysregulated REs at locus-specific levels in multiple cancer types. With dysregulation in any seven cancer types analyzed, we defined the recurrently differentially expressed REs (see **Fig. 2c and 2d**). Interestingly, genes corresponding to the recurrently up-regulated genic REs are enriched in DNA replication and mitotic cell cycle. In contrast, genes corresponding to the recurrently down-regulated genic REs are enriched in the extracellular matrix. Notably, among these genes, 12 cancer-related genes, including CDK1 (oncogene^50^), ESCO2 (oncogene^51^), GAS5 (tumor suppressor non-coding gene^52^), GTSE1 (oncogene^53^), H2BC12 (potential oncogene^54^), KIFC1 (oncogene^55^), MMS22L (oncogene^56^), NF1 (tumor suppressor gene^57^), NIT2 (tumor suppressor gene^58^), PVT1 (non-long coding RNA with oncogenic effects^59^), STAT1 (tumor suppressor gene^60^) and UBE2C (oncogene^61^) are consistently up-regulated (see **Fig. S6a**). At the same time, 17 out of 19 cancer-related genes, including ADAMTS9-AS2 (tumor suppressor long non-coding gene^62^), ATF3 (tumor suppressor gene^63^), DCN (tumor suppressor gene^64^), DUSP1 (promote carcinogenesis in some cancers and inhibits carcinogenesis in other cancers^65^), ECT2L (oncogene^66^), EMP1 (oncogene^67^), JDP2 (tumor suppressor gene^68^), KLF6 (tumor suppressor gene^69^), NDRG2 (tumor suppressor gene^70^), NR4A1 (tumor suppressor in some cancers oncogene in other cancers^71^), RHOBTB2 (tumor suppressor gene^72^), SOCS2 (tumor suppressor gene^73^), SPARCL1 (tumor suppressor gene^74^), STARD13 (tumor suppressor gene^75^), SYNPO2 (tumor suppressor gene^76^), TAGLN (oncogene^77^), and TIMP3 (tumor suppressor gene^78^) are mostly down-regulated in most of the 12 cancer types (see **Fig. S6b**). Furthermore, since one gene typically consists of many transcripts and each transcript can be associated with many genic REs, we found the regulation direction (*i.e.*, up or down-regulation) for a given gene is similar to the regulation direction of the majority of the corresponding genic REs as shown in **Fig. S6c-S6f**.

Among five consistently down-regulated genic REs in all 12 cancer types, four are in the same intronic region of gene TMEM252 (with transcripts ID: NM_153237.2), as shown in **Fig. 6a**. Interestingly, one of these fifth RE elements (*i.e.*, LTR82A_dup582) is also located in the same intronic region, however, with little to no read’s coverage in almost all 12 cancer types (see **Fig. 6b, 6c** and **Fig. S8**), indicating the less likely event of read-through transcription. TMEM252 (Human transmembrane protein 252), a member of the transmembrane protein family, showed significantly reduced expression in the majority of 12 cancer types (including BLCA, BRCA, COAD, ESCA, HNSC, LIHC, LUAD, PRAD, THCA) with non-significant reductions in UCEC comparing tumor with matched normal samples as shown in **Fig. S8**. Interestingly, even with all four corresponding genic REs being significantly down-regulated in KIRC and KIRP, the expression level of TMEM252 showed comparable expression between the tumor and matched normal samples (see **Fig. S8**). TMEM252 has recently been identified as a tumor suppressor gene in triple-negative breast cancer to inhibit its progression by suppressing STAT3 activation^79^. Furthermore, a recent study on papillary thyroid carcinoma demonstrated that overexpression of TMEM252 can suppress cell proliferation by repressing the expression of p53, p21, and p16 through the inhibition of the Notch pathway and consequently epithelial-mesenchymal transition, overexpression of TMEM252 also inhibited the cell migration and invasion^80^. Although not being shown down-regulated in KIRC and KIRP in this study, evidence showed that the higher expression level of TMEM252 in KIRC (https://www.proteinatlas.org/ENSG00000181778-TMEM252/pathology/renal+cancer/KIRC) and KIRP (https://www.proteinatlas.org/ENSG00000181778-TMEM252/pathology/renal+cancer/KIRP) are associated with a higher survival probability for the cancer patients. Given the consistent down-regulation across the vast majority of 12 cancer types analyzed in this study and the recently reported tumor-suppressing effects, TMEM252 could potentially act as a tumor suppressor gene in a wide range of cancer types, thus warranting further in-depth investigation.

To investigate the epigenetic dysregulations of REs, we also analyzed the DNA methylation changes of REs that can be uniquely identified in their 1kb TSS by the corresponding methylation probes. Compared with RNA-seq data, only about 1.5% of REs (66,859 out of 4,467,488) are available for analysis of methylation changes due to the low coverage of Illumina 450K methylation array in the human genome (*i.e.*, only 1.5% CpG coverage in the human genome^39^). In contrast to differentially expressed REs (see **Table S2**), 10 out of 12 cancer types (BLCA, BRCA, COAD, ESCA, HNSC, LIHC, LUAD, THCA, and UCEC) showed a higher number of hypo-methylated REs at locus-specific levels than hyper-methylated ones (see **Table S17**). This is consistent with the TE methylation changes observed at the subfamily level^3^, further indicating the validity of the RE methylation analysis conducted in this study at the locus-specific level.

Despite the general belief that DNA methylation negatively regulates RE expression^1,12^, our analysis revealed a discrepancy between the prevalence of hypo-methylated REs and up-regulated REs in cancer types: namely, the larger number of cancer types with higher levels of hypo-methylated REs and the smaller number of cancer types showing up-regulated REs. This discrepancy may be partially attributed to the limited coverage of the Illumina 450K methylation array. To overcome the small overlaps between differentially methylated REs and differentially expressed REs shown in **Fig. S11** and to explore the association between RE methylation and expression changes, we compared the expression changes between hypo- and hyper-methylated REs as well the averaged methylation changes between up- and down-regulated REs shown in **Fig. 7a** and **7b** respectively. We also checked the correlation between methylation and expression changes for recurrently dysregulated REs covered by the DNA methylation array (see **Fig. 7c** and **7d**). In general, expression changes are higher for hypo-methylated REs than hyper-methylated ones, and averaged methylation changes are lower for up-regulated REs than down-regulated ones. Although the Illumina 450K methylation array’s limited coverage restricted our analysis of the association between RE methylation and expression, our results are broadly consistent with the belief that DNA methylations restrict RE activities.

To our knowledge, this is the first study that comprehensively characterized the dysregulation of locus-specific REs among multiple common cancer types. With the increasing adaptation to long-read sequencing^81^, we expect a higher interest in studying REs at locus-specific levels in cancer research. Due to the limited coverage of the Illumina 450K methylation array, a comprehensive understanding of methylation changes for REs at locus-specific levels in the common cancer is still lacking. With the increasing application of whole genome bisulfite sequencing in cancer studies^82^, we expect a high-resolution map of DNA methylation changes for different cancer types. Future studies utilizing whole-genome bisulfite sequencing will provide a more comprehensive picture of DNA methylation patterns for REs at the locus-specific level. Considering the rich information contained in REs at the locus-specific level and complementary information offered by RE expression and methylation changes, we expect to see the development of RE-based biomarkers for cancer type classification and, eventually, for potential therapeutic targets in the near future.

