## Supplementary Figures for "Dysregulation of locus-specific repetitive elements in TCGA pan-cancers"

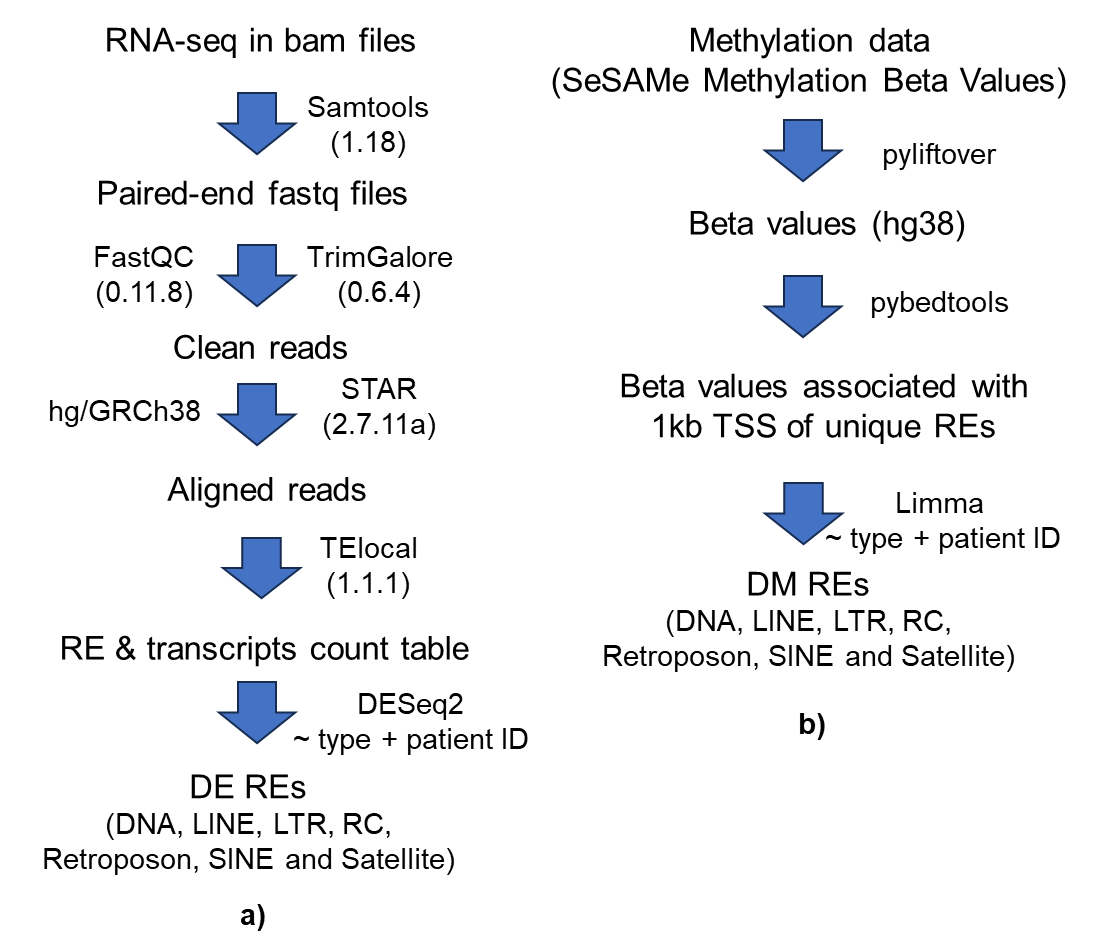

**Fig. S1:** The workflow to identify dysregulated REs across 12 cancer types. **a):** The workflow for RE expression analysis; **b):** The workflow for RE methylation analysis.

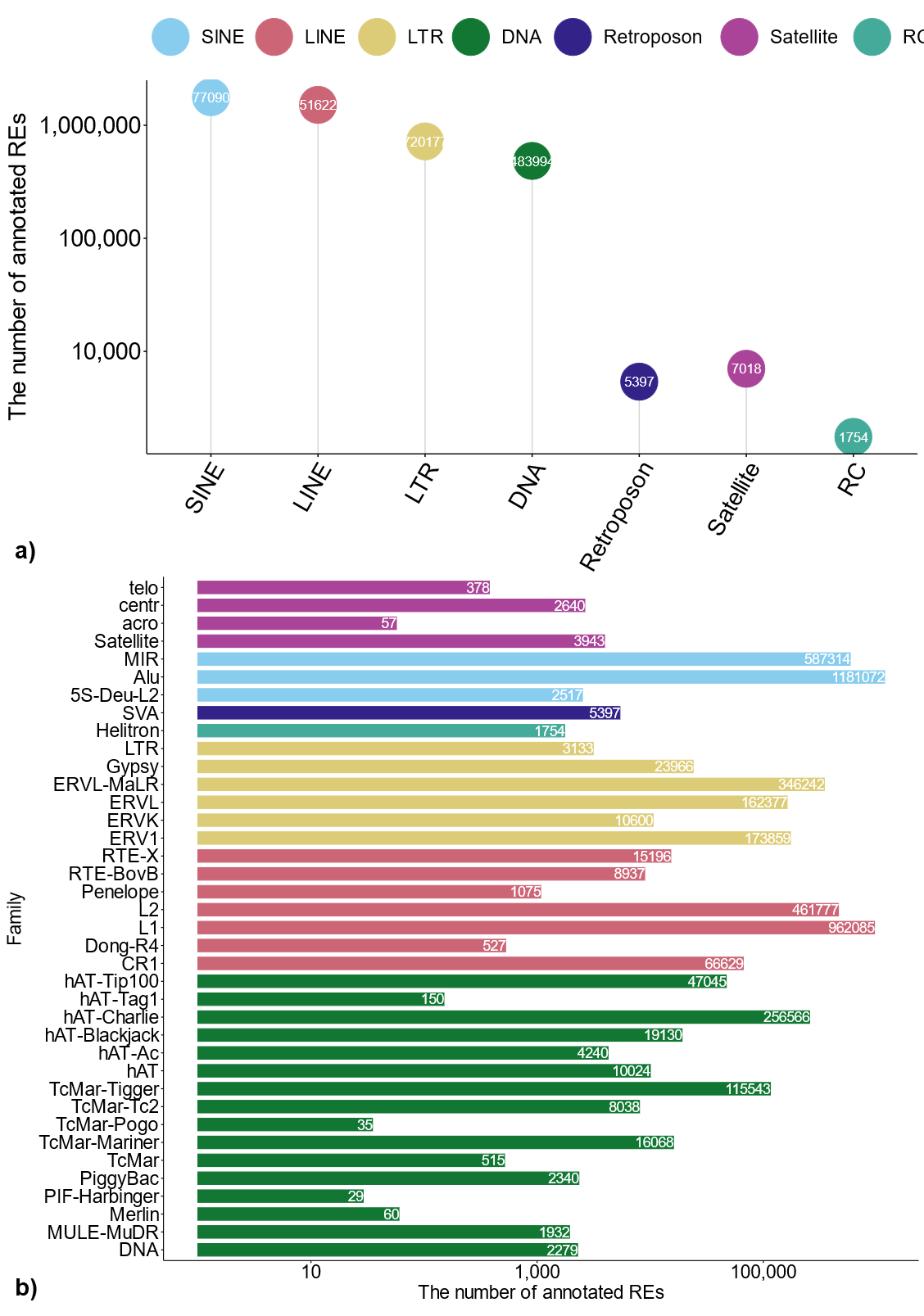

**Fig. S2: Summary of locus-specific REs based on hg38 annotation for TElocal. a):** Number of locus-specific RE elements for each of 7 RE classes; **b):** Number of locus-specific RE elements in each RE family.

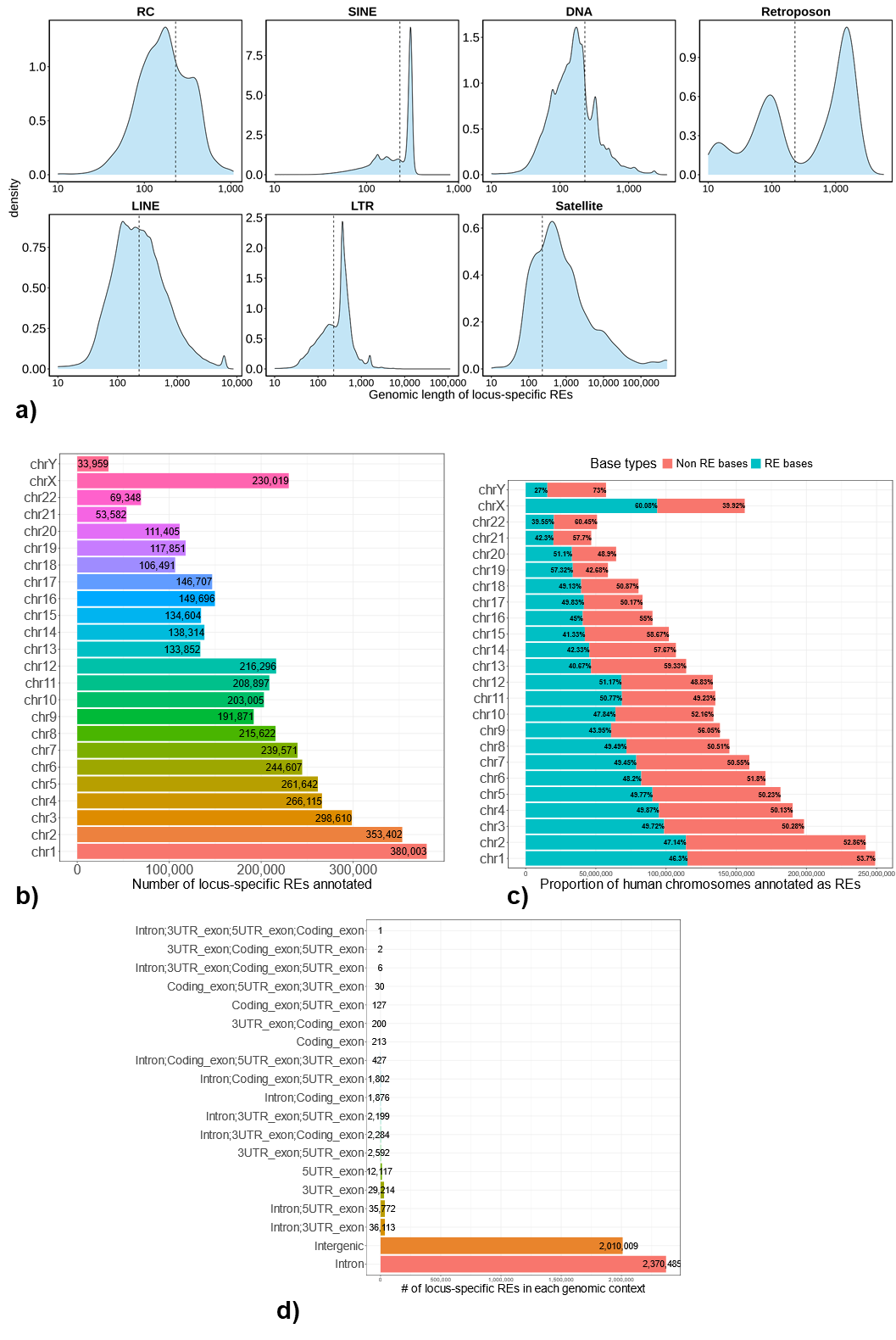

**Fig. S3: Distribution of locus-specific REs in the human genome based on hg38 annotation for TElocal. a):** genomic length of each RE class in the human genome; **b):** number of locus-specific REs in each chromosome; **c):** percentage of nucleotide bases in each chromosome that are annotated as REs; **d):** number of locus-specific REs with different genomic contexts based on protein-coding genes.

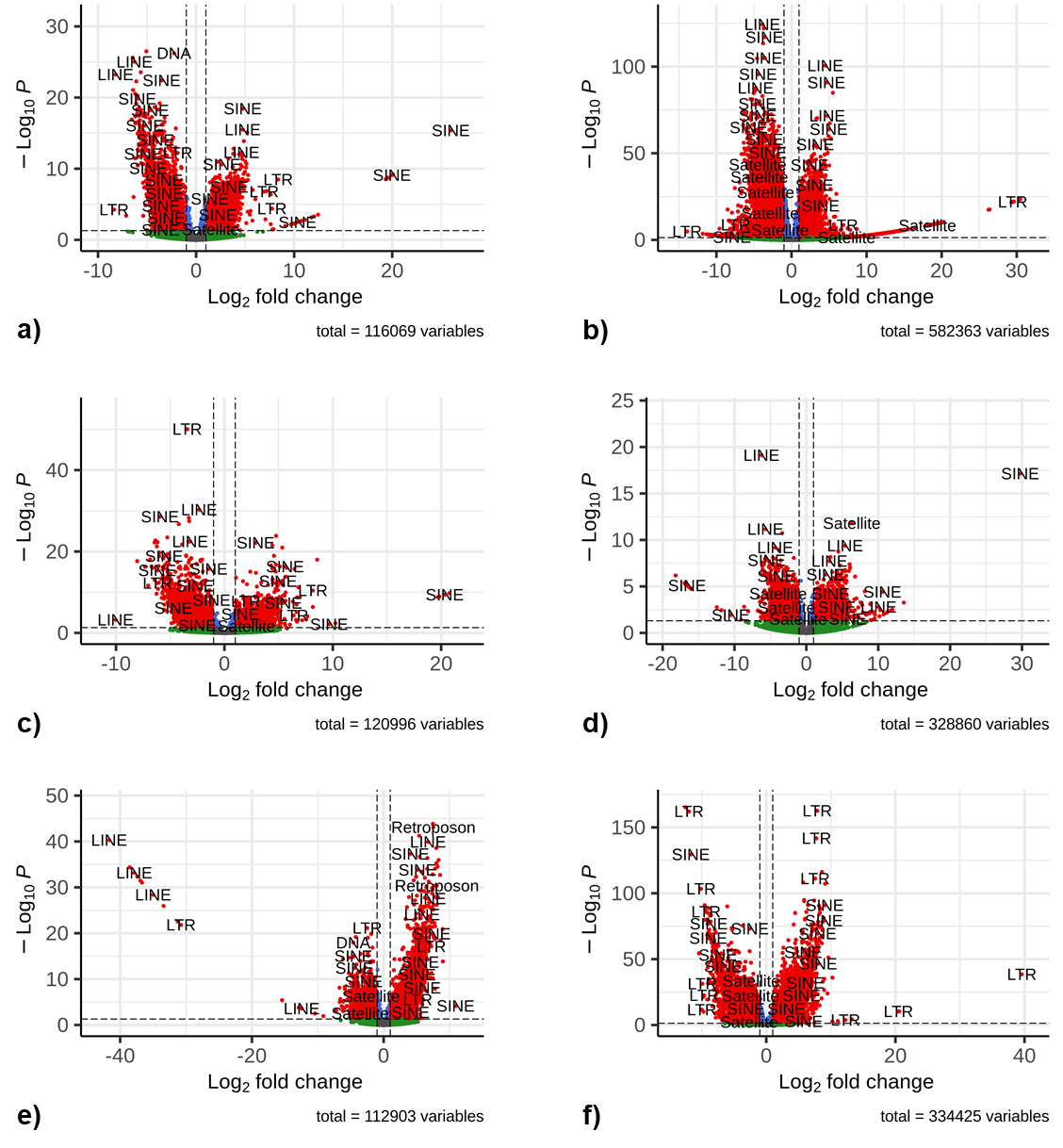

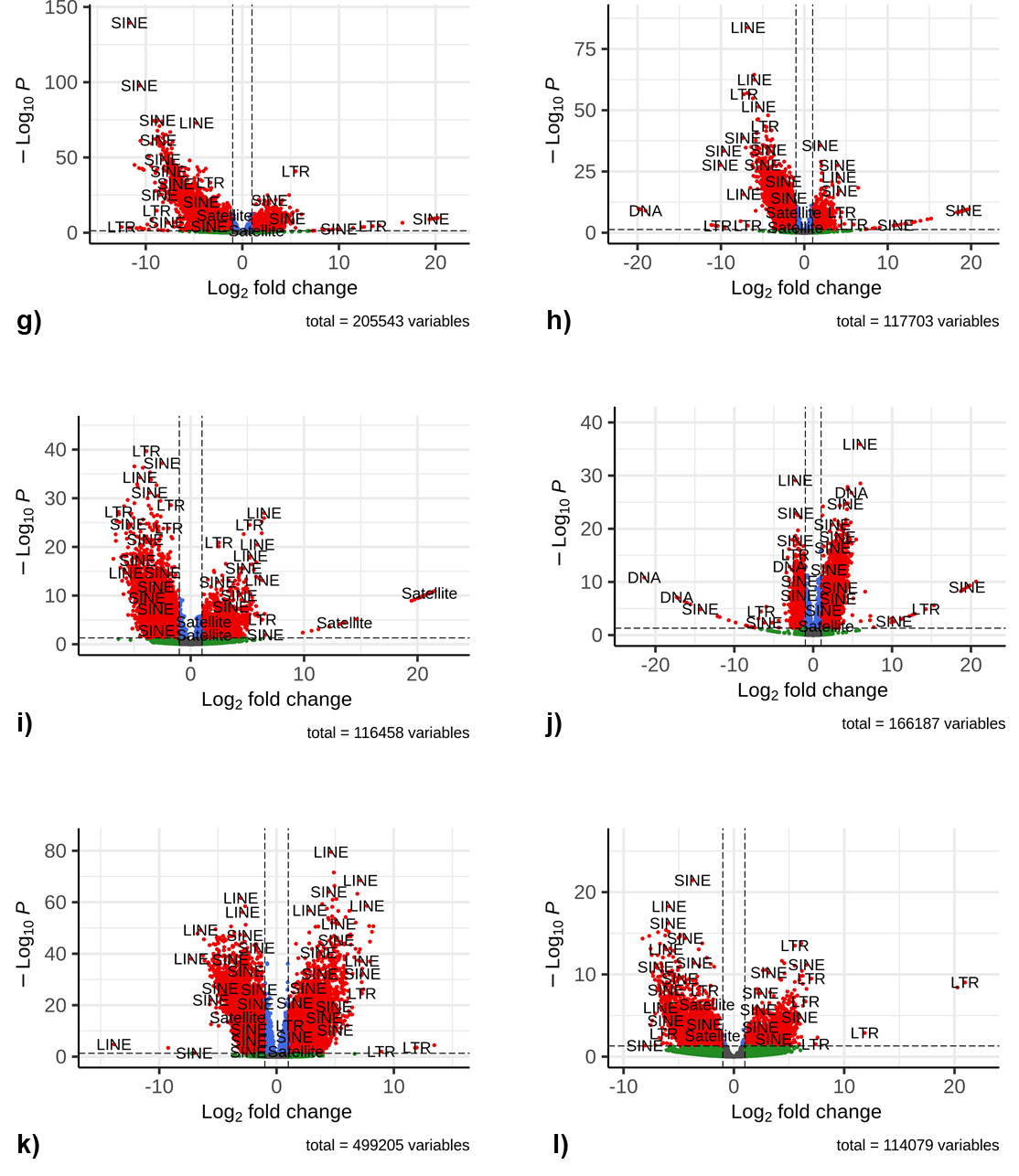

**Fig. S4: Expression changes of locus-specific REs in each of the 12 cancer types** (REs with |log2 fold change| >= 1 and adjusted p-values <= 0.05 are considered differentially expressed REs and represented by red points)**. a):** expression changes of locus-specific REs for BLCA between tumors and matched normal samples; **b):** for BRCA; **c):** for COAD; **d):** for ESCA; **e):** for HNSC; **f):** for KIRC; **g):** for KIRP; **h):** for LIHC; **i):** for LUAD; **j):** for PRAD; **k):** for THCA; **l):** for UCEC;

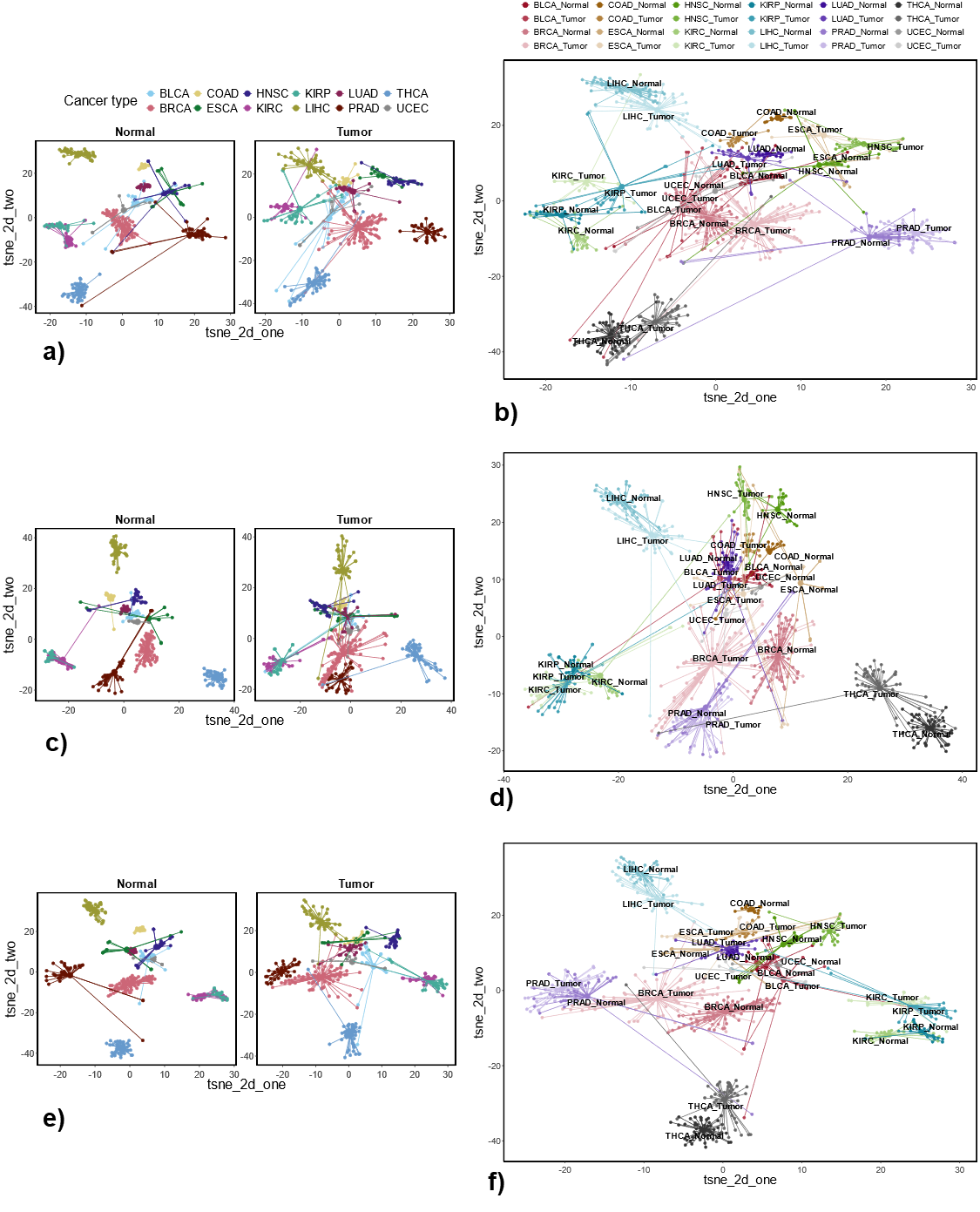

**Fig. S5. Sample clustering with uniquely dysregulated intergenic REs identified across 12 cancer types. a):** t-SNE plots based on uniquely up-regulated intergenic REs for Normal and Tumor sample clustering separately; **b):** t-SNE plot based on uniquely up-regulated intergenic REs for different sample type clustering (tumor with matched normal samples); **c):** t-SNE plots based on uniquely down-regulated intergenic REs for Normal and Tumor sample clustering separately; **d):** t-SNE plot based on uniquely down-regulated intergenic REs for different sample type clustering (tumor with matched normal samples); **e):** t-SNE plots based on uniquely up- and down-regulated intergenic REs for Normal and Tumor sample clustering separately; **f):** t-SNE plot based on uniquely up- and down-regulated intergenic REs for different sample type clustering (tumor with matched normal samples);

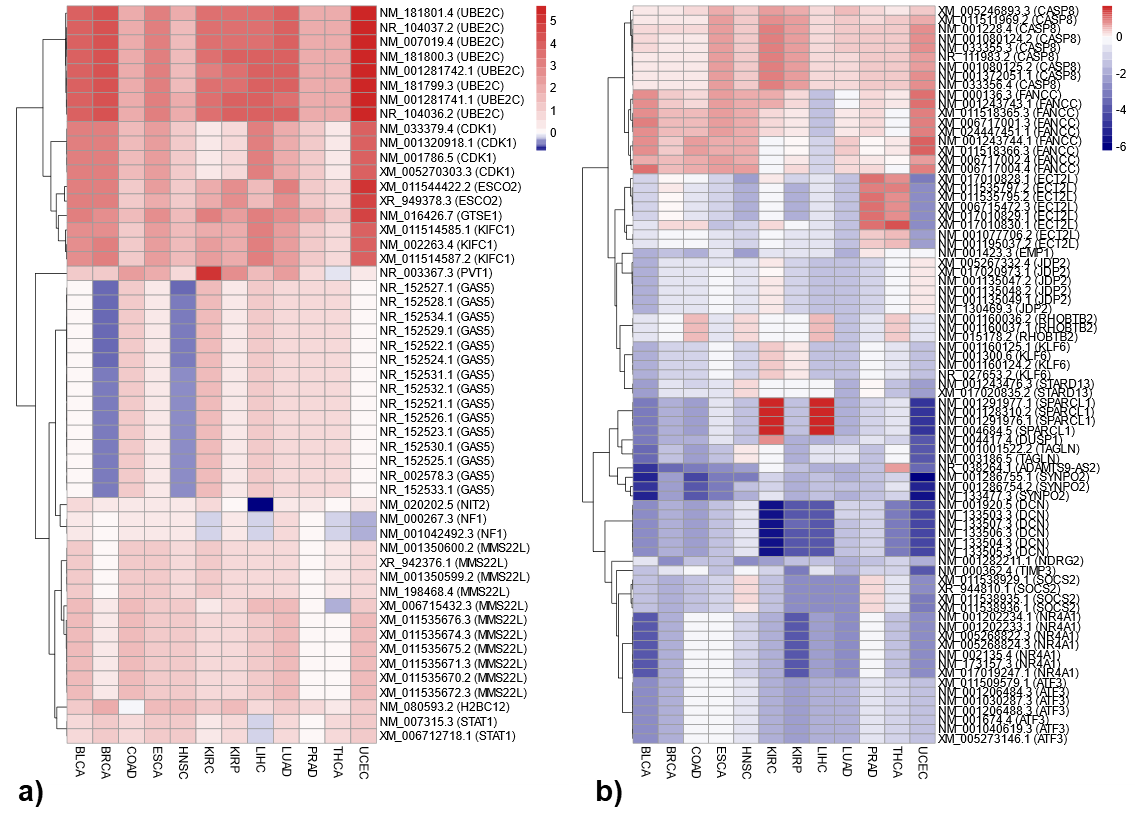

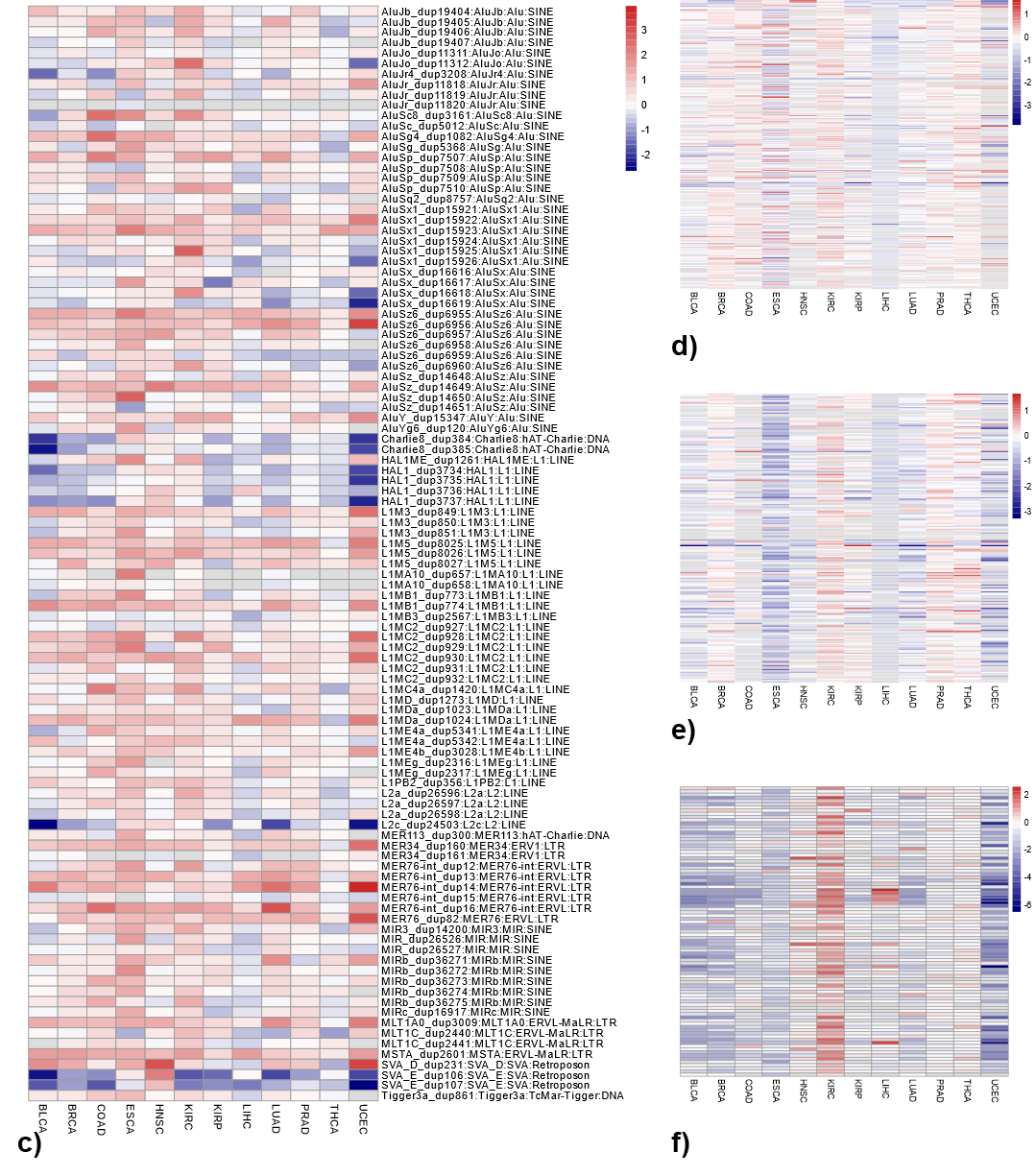

**Fig. S6: Expression changes of transcripts corresponding to cancer genes associated with recurrently dysregulated genic REs. a):** expression changes of transcripts (that are cancer genes) associated with recurrently up-regulated genic REs; **b):** expression changes of transcripts (that are cancer genes) associated with recurrently down-regulated genic REs; **c):** expression changes of REs that are associated with gene CASP8; **d):** expression changes of REs that are associated with gene FANCC; **e):** expression changes of REs that are associated with gene ECT2L; **f):** expression changes of REs that are associated with gene SPARCL1;

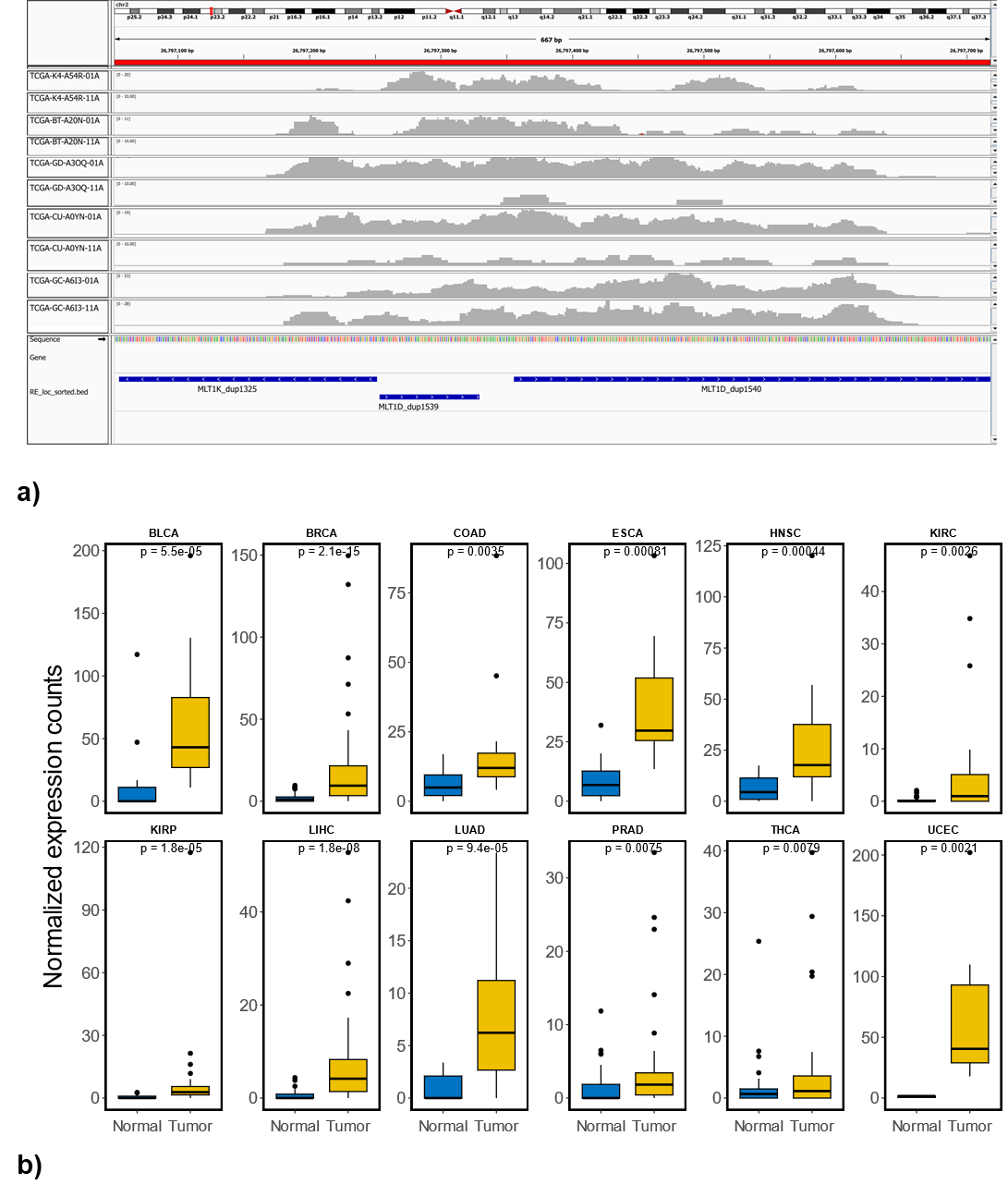

**Fig. S7: Consistent up-regulated intergenic RE** (*i.e.*, MLT1D_dup1540) **across 12 cancer types. a):** reads coverage corresponding to MLT1D_dup1540 between the randomly selected 5 tumor and matched normal samples in BLCA (tumor sample ends with 01A while the matched normal sample ends with 11A); **b):** normalized expression comparison between tumor and matched normal samples for MLT1D_dup1540 across 12 cancer types.

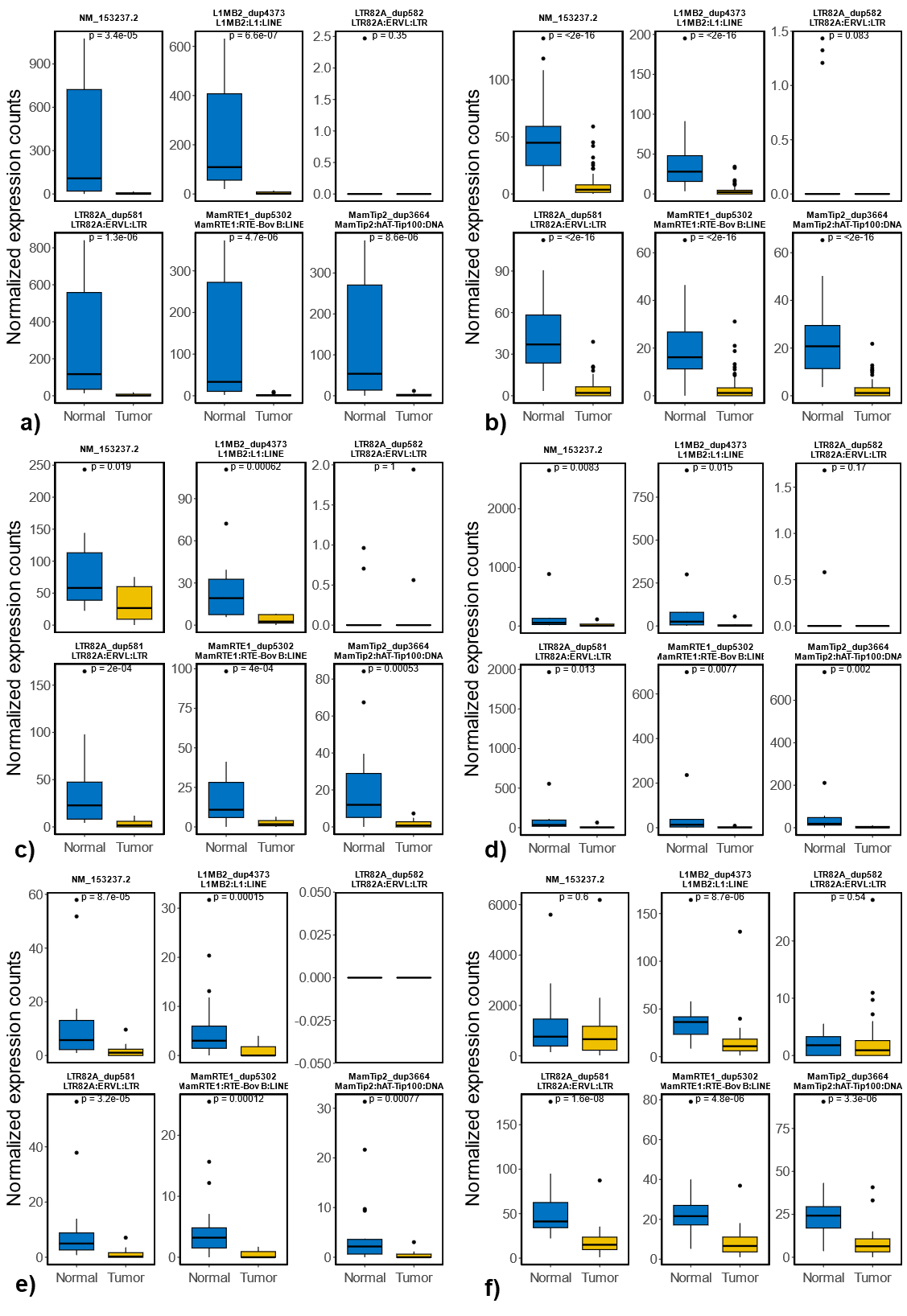

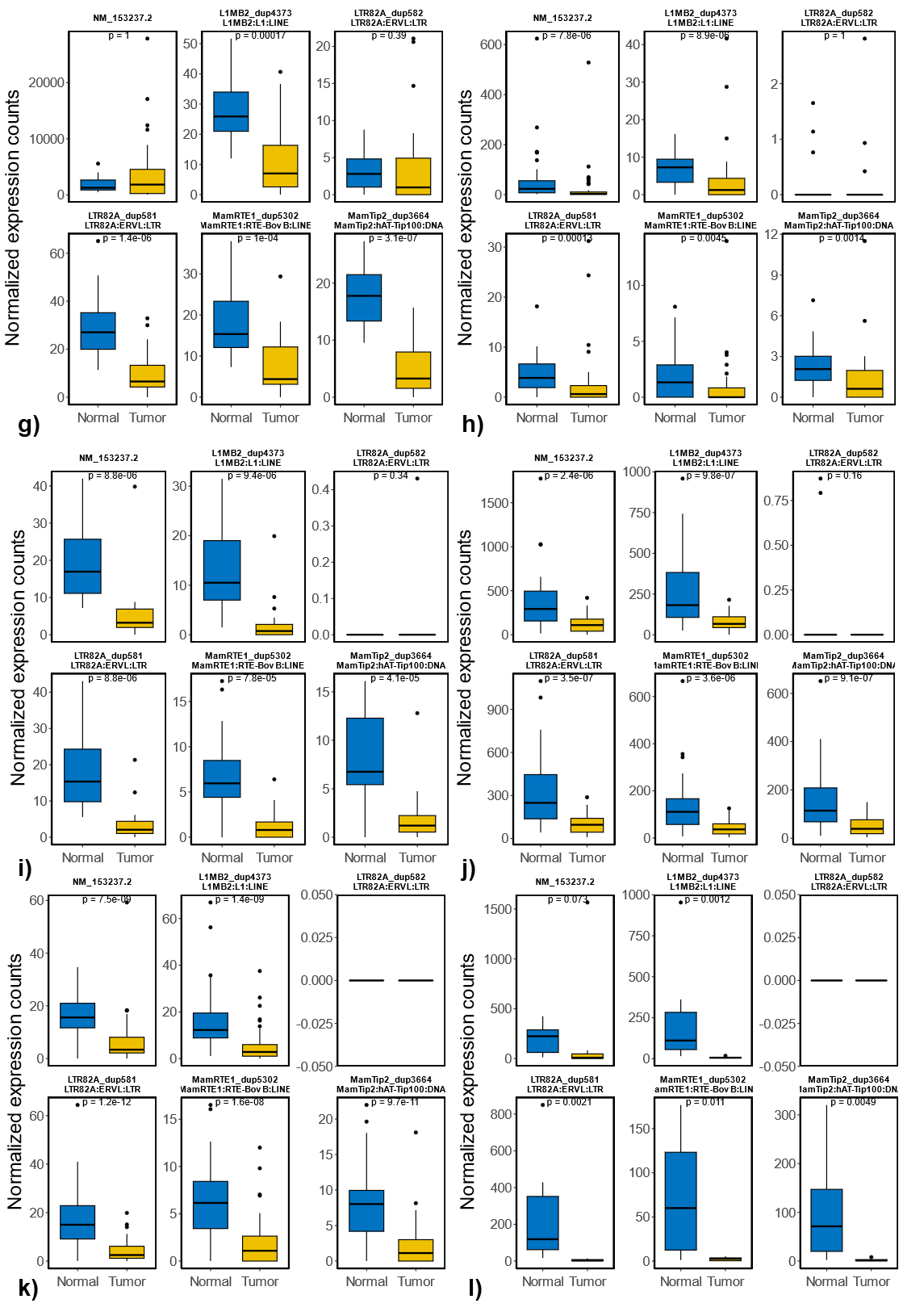

**Fig. S8: Expression comparison between tumor and matched normal samples for TMEM252 gene (with transcripts ID: NM_153237.2) and its associated REs across 12 cancer types.** **a):** for BLCA; **b):** for BRCA; **c):** for COAD; **d):** for ESCA; **e):** for HNSC; **f):** for KIRC; **g):** for KIRP; **h):** for LIHC; **i):** for LUAD; **j):** for PRAD; **k):** for THCA; **l):** for UCEC;

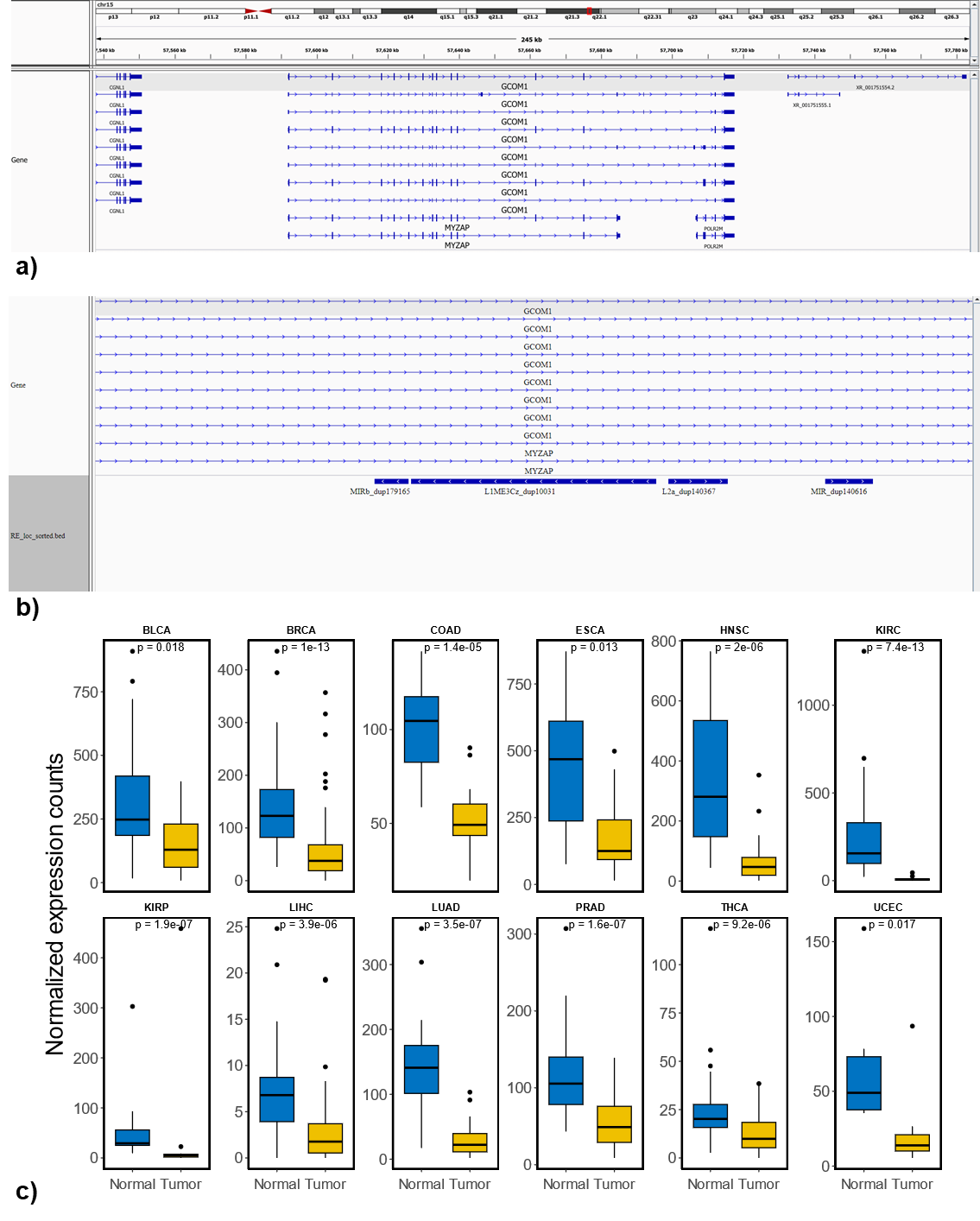

**Fig. S9: Consistent down-regulated intronic RE** (*i.e.*, L1ME3Cz_dup10031) **across 12 cancer types. a): annotation for** L1ME3Cz_dup10031 associated genes; **b):** genomic context corresponding to L1ME3Cz_dup10031 in the human genome; **c):** normalized expression comparison between tumor and matched normal samples for L1ME3Cz_dup10031 across 12 cancer types.

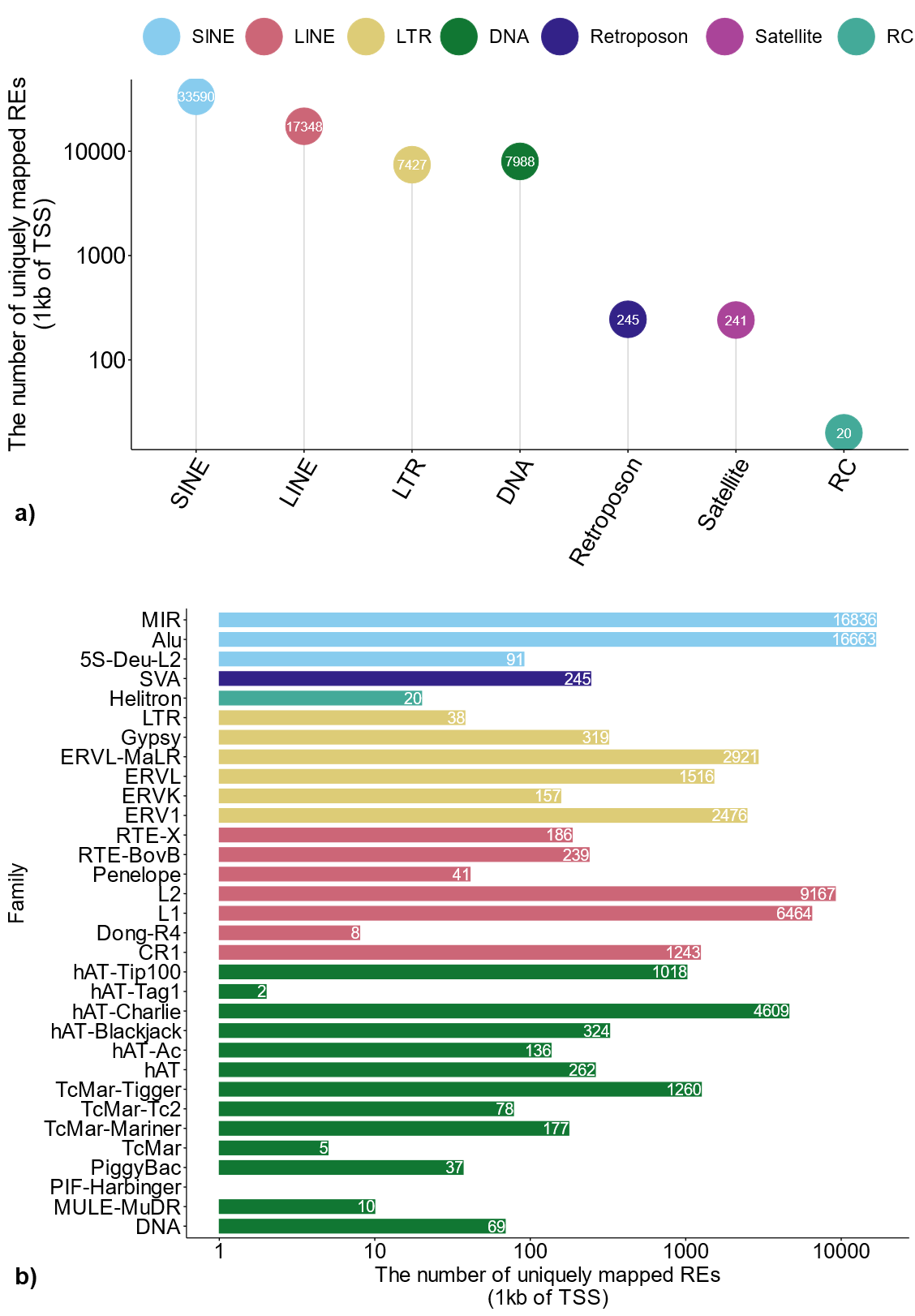

**Fig. S10: Number of REs that can be uniquely mapped with DNA methylation probes. a):** Number of locus-specific RE elements for each of 7 RE classes that can be uniquely mapped with DNA methylation probes; **b):** Number of locus-specific RE elements in each RE family that can be uniquely mapped with DNA methylation probes.

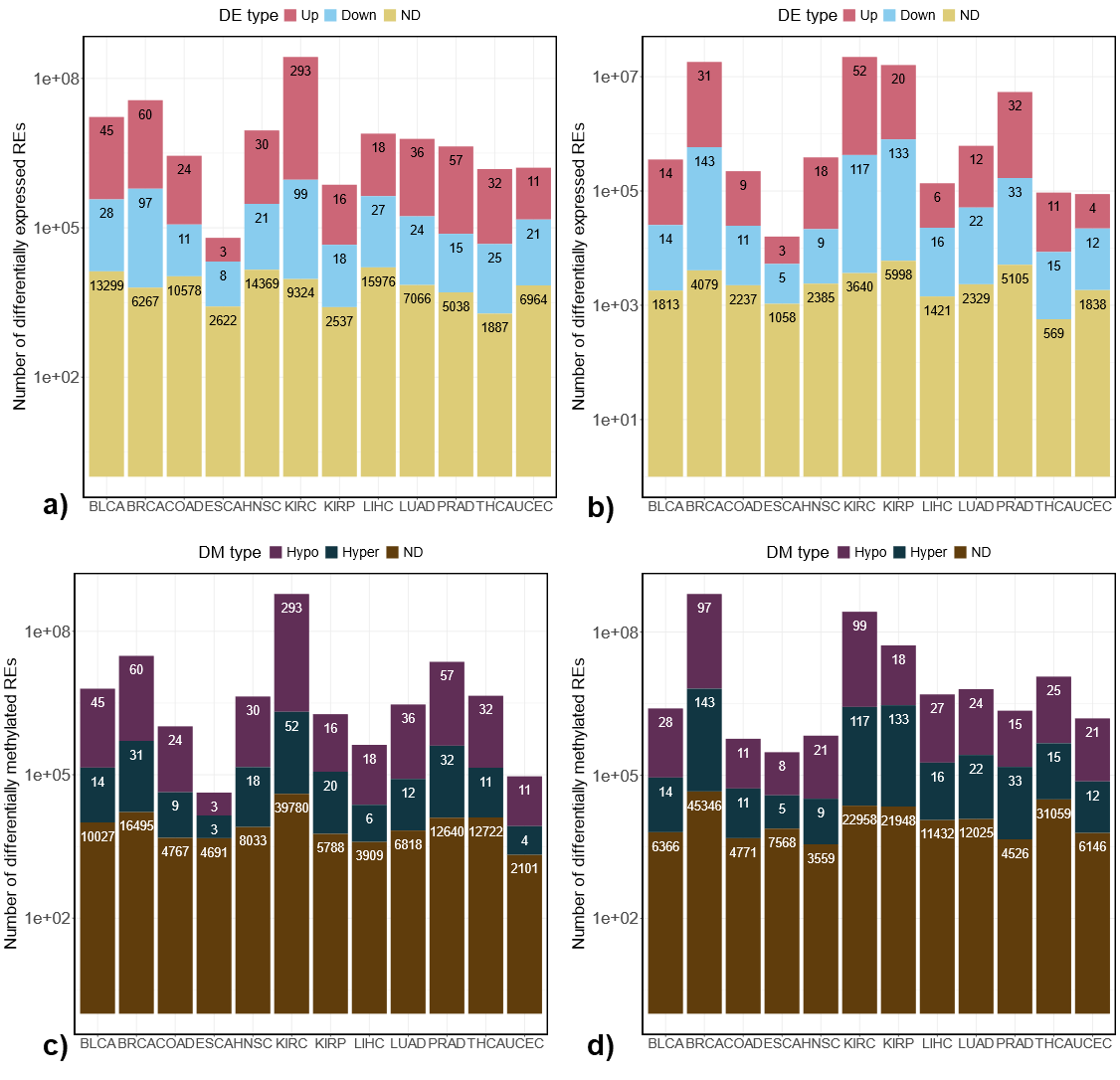

**Fig. S11: Number of REs that are differentially expressed and differentially methylated** (Up: up-regulated; Down: down-regulated; ND: no significant difference; Hypo: hypo-methylated; Hyper: hyper-methylated). **a):** number of differentially expressed REs that are hypo-methylated across all 12 cancer types; **b):** number of differentially expressed REs that are hyper-methylated across all 12 cancer types; **c):** number of differentially methylated REs that are up-regulated across all 12 cancer types; **d):** number of differentially methylated REs that are down-regulated across all 12 cancer types.
